# Delineating markers of disease-disease interaction: a systematic methodology and its application to multiple diabetes-helminth cohorts

**DOI:** 10.1101/2023.12.27.573481

**Authors:** Nilesh Subramanian, Philge Philip, Anuradha Rajamanickam, Nathella Pavan Kumar, Subash Babu, Manikandan Narayanan

**Affiliations:** Department of Computer Science and Engineering, Indian Institute of Technology (IIT) Madras, Chennai, India; Center for Integrative Biology and Systems Medicine, IIT Madras, Chennai, India; NIH-NIAID-International Center for Excellence in Research, Chennai, India; ICMR-National Institute for Research in Tuberculosis, Chennai, India; Helminth Immunology Section, Laboratory of Parasitic Diseases, NIAID-NIH, Bethesda, MD, USA; Wadhwani School of Data Science and Artificial Intelligence, IIT Madras, Chennai, India

## Abstract

Understanding how the molecules in our body respond to the co-occurrence of two diseases in an individual (comorbidity) could lead to mechanistic insights into novel treatments for comorbid conditions. Studies have shown for instance that responses of our immune system to comorbid conditions could be more complex than the union of immune responses to each disease occurring separately, but a data-driven quantification of this complexity is lacking. In this study, we present a systematic methodology to quantify the interaction effect of two diseases on marker variables of interest (using a chronic inflammatory disease diabetes and parasitic infection helminth as illustrative disease pairs to identify cytokines or other immune markers that respond distinctively under a comorbid condition). To perform this systematic comorbidity analysis, we (i) collected and preprocessed data measurements from multiple single- and double-disease cohorts, (ii) extended differential expression analysis of such data to identify disease-disease interaction (DDI) markers (such as cytokines that respond antagonistically or synergistically to the double-disease condition relative to single-disease states), and (iii) interpreted the resulting DDI markers in the context of prior cytokine/immune-cell knowledgebases.

We applied this three-step DDI methodology to multiple cohorts of helminth and diabetes (specifically, helminth-infected and helminth-treated individuals in diabetic and non-diabetic conditions, and non-disease control individuals), and identified cytokines such as IFN-*γ*, TNF-*α*, and IL-2 to be DDI markers acting at the interface of both diseases in data collected prior to helminth treatment. Validating our expectations, for these cytokines and other T helper Th-2 cytokines like IL-13 and IL-4, their DDI statuses were lost after treatment for helminth infection. For instance, the relative contribution of the DDI term in explaining the individual-to-individual variation of IFN-*γ* and TNF-*α* cytokines were 67.68% and 48.88% respectively before anthelmintics treatment and dropped to 6.09% and 14.56% respectively after treatment. Furthermore, signaling pathways like IL-10 and IL-4/IL-13 were found to be significantly enriched for genes targeted by certain DDI markers, thereby suggesting mechanistic hypotheses on how these DDI markers influence both diseases. Our results quantified the extent of helminth-diabetes DDI exhibited by various tested cytokine markers, and thereby delineated their role in the pathogenesis of both diseases. These results are promising and encourage the application of our DDI methodology (https://github.com/BIRDSgroup/DDI) to dissect the interaction between any two diseases, provided multi-cohort measurements of markers are available.

## Introduction

Comorbidity refers to the condition when more than one disease affects an individual at the same time. For instance, an acute infectious disease can afflict an individual suffering from a chronic disease (such as diabetes), and such comorbid conditions can lead to an interesting effect on the various immune/inflammatory molecules released by the host system. For instance, the effect on immune molecules when two diseases interact within an individual could be quite different (synergistic or antagonistic say, rather than simply being additive) compared to when each disease affects the individual in isolation. Quantifying the distinctive double-disease effect on molecules of interest can yield insights into the dynamics of disease progression under comorbid conditions and help reveal the biological pathways affected by these molecules. Genes in such biological pathways can be experimentally validated for their links to the two diseases, and subsequently targeted to design novel comorbidity therapies. Note that a comorbidity therapy may offer advantages that go beyond a simple combination of the individual treatments for each disease.

Since studying the pathophysiology of any disease in isolation is challenging in and of itself, it is even more challenging to dissect the pathophysiology of a comorbid condition [1]. Some of the challenges in identifying key players involved in comorbid conditions include the lack of carefully planned cohorts comprising single- vs. double-disease cases and appropriate controls, low sample sizes of such multi-cohort datasets even if available, and statistical challenges like multiple testing burden and correction for confounding covariates. While certain studies have identified comorbidity molecules from multi-cohort datasets [2], a rigorous computational approach to quantify the double-disease effect on such molecules and interpreting them biologically is still lacking. In this study, we take a systematic approach to quantify the interaction effect of two diseases on marker variables or molecules of interest (using diabetes and helminth as illustrative disease pairs to identify cytokines or other immune markers that respond distinctively under a comorbid condition). More specifically, we propose a three-step computational pipeline wherein the single and double disease contributions to each marker variable are estimated (using a linear regression model with main and interaction effects [3–5], which extends a standard case vs. controls differential expression analysis), to identify markers that exhibit statistically significant disease-disease interaction (DDI) effect, and interpret the resulting DDI markers in the context of prior immune-cell or cytokine knowledge-bases.

The application of our DDI pipeline to the two diseases, diabetes (which refers to type 2 diabetes mellitus in this work) and helminth (specifically parasite *Strongyloides stercoralis* or *Ss* infection), is motivated by epidemiological and laboratory studies that have shown helminth infections to be associated with a decreased prevalence of diabetes [6]. This inverse helminth-diabetes association is one aspect of the hygiene hypothesis [7, 8], and can be understood as follows.

(disease-1 response): Diabetes is a well-known metabolic disease associated with a wide range of inflammatory immune responses involving T-helper cells Th1 and Th17 [9, 10]. Particularly inflammatory cells like neutrophils and lymphocytes are said to produce high levels of chemical molecules, causing high inflammation [11].
(disease-2 response): Helminth infection is characterized by profound secretion of Th2 cytokine responses and regulatory cytokines which may contribute to the overall protective response to these infections [12–14].
(interaction response): Therefore, in the presence of helminth infections, the immune system is in an anti-inflammatory mode deemed disadvantageous to diabetes progression. Berbudi et al. in their work illustrate a possible mechanism of an inverse relationship between helminth infections and diabetes by saying that type-2 immune response and its associated cell types like alternatively activated macrophages are activated by helminth infections thereby protecting against diabetes [15]. In their research, Camaya et al. speculate that the inverse link between helminth infections and diabetes may be due to a potential cross-talk between the macrophages activated by helminth infection and *β*-cells of the pancreas. [16]. Differential analysis of samples with diabetes and with/without the helminth infection, carried out by Rajamanickam et al. [2], showed that compared to uninfected individuals, infected individuals have significantly diminished levels of insulin, glucagon, adiponectin, adipsin, Th17, and Th1 cytokines, and these levels reversed following anthelmintic therapy.

While some information as indicated above is known about the mechanisms of interaction between the two diseases, what is lacking is a quantitative scoring and systematic screening of immune cells and inflammatory molecules that behave distinctively under the comorbid helminth-diabetes condition.

In this study, we apply our proposed DDI pipeline to perform interaction analysis of immune or inflammatory markers measured in four groups of individuals: control, disease-1, disease-2, and comorbid individuals. We apply it to investigate cellular/molecular markers at the interface of diabetes and helminth infection. More specifically, this pipeline allowed us to estimate the per-disease main effects and DDI (interaction) effect on each measured marker. Thus, we could identify and interpret statistically significant DDI markers (cytokines) such as IFN-*γ*, TNF-*α*, and IL-2 and the loss of their DDI statuses after anthelmintic treatment. For instance, the relative contribution of the helminth, diabetes, and interaction terms to explaining the variance of the DDI marker IFN-*γ* are 32.1%, 0.2%, and 67.7% in the before-treatment cohorts, and shift to 21.6%, 72.3%, and 6.1% in the after-treatment cohorts. Various pathways like IL-10, IL-4, and IL-13 signaling are seen to be enriched for genes that are targeted by the DDI markers, suggesting mechanisms through which the two diseases interact. Our study has promising applications beyond helminth and diabetes as well since our DDI methodology applied on single- and double-disease cohorts data can reveal the impact of comorbidity on any measured marker for any disease pair of interest, thereby allowing screening for molecules that lie at the interface of the two diseases.

## Results

### Our Multi-cohort Disease-Disease Interaction (DDI) Analysis Overview

For any two diseases of interest, a biochemical variable (such as a cytokine’s concentration level or immune celltype’s frequency) is referred to as a Disease-Disease Interaction (DDI) marker if the change in this variable in a group of comorbid individuals relative to the controls group is significantly different from the cumulative changes in the single-disease groups (i.e., the sum of changes of this variable in disease 1 vs. controls and disease 2 vs. controls). Our DDI pipeline identifies such DDI markers by performing linear regression-based interaction analysis of variables measured in a carefully curated set of single-disease, double-disease, and control groups/cohorts of individuals (see Fig 1); and provides additional interpretations of the identified DDI markers by querying them in prior information sources (e.g., cytokine-gene knowledgebases). This overall structured pipeline, comprising the three steps of preprocessing of cohorts, differential expression analysis involving main (per-disease) and interaction (DDI) effect terms, and downstream interpretation of the identified markers in the context of prior biological knowledge, is depicted in Fig 1A and described in detail in Methods.

**Fig 1.**
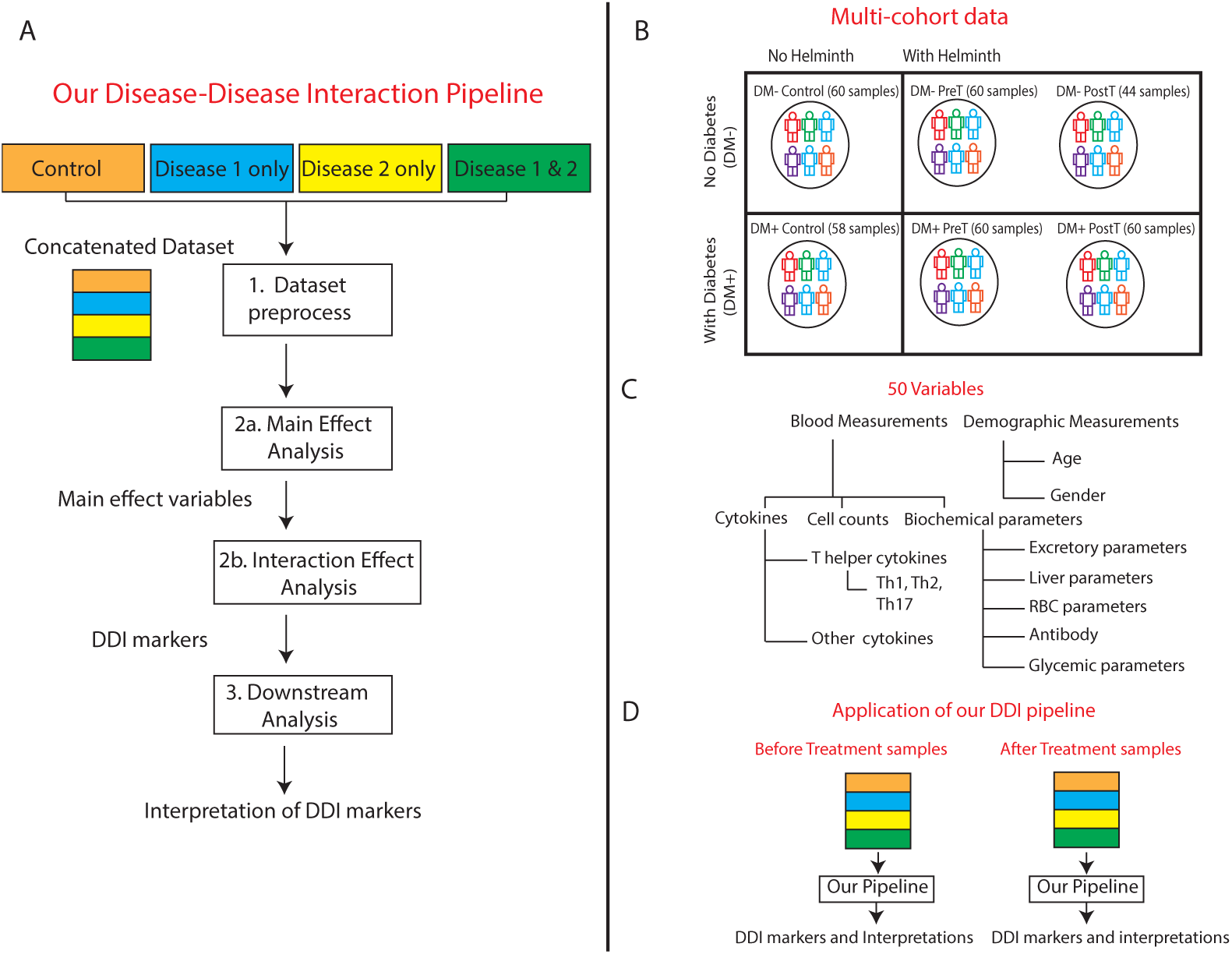
Overview of our DDI methodology and dataset: (A) Our DDI pipeline analyzes variables measured in multiple cohorts (control, disease 1, disease 2, and both diseases together) to reveal the double-disease interaction markers. (B) Multiple cohorts of diabetes and helminth infection (both pre- and post-anthelmintic treatment, denoted respectively as PreT and PostT) on which our DDI pipeline is applied. (C) The different blood-based and demographic variables measured in each cohort (see also Suppl File D1; RBC stands for Red Blood Cell). (D) In the current work, our pipeline was applied to two sets of diabetes-helminth cohorts, namely before-treatment cohorts (DM*−* Control, DM*−* PreT, DM+ Control, DM+ PreT) and after-treatment cohorts (DM*−* Control, DM*−* PostT, DM+ Control, DM+ PostT). Treatment in this paper refers to treatment for helminth.

In the application of our DDI pipeline to study helminth-diabetes comorbidity, we used published multi-cohort data [2] gathered from 118 diabetes mellitus individuals, including 60 helminth-infected people (DM+ Pre-treatment) and 58 people who had no infection (DM+ Control). Data without diabetes was gathered from 120 people, including 60 without helminth infection (DM*−* Control) and 60 with helminth infection (DM*−* Pre-Treatment). In both datasets, anthelmintic treatment was administered to all infected individuals (DM+ Post-treatment, DM*−* Post-treatment), and measurements were repeated six months later (except for a subset of individuals who couldn’t be followed up for measurements in DM*−* Post-treatment; see Methods). That is, demographic and blood-based measurements were taken from individuals in both pre- and post-helminth treatment time points (referred to as pre-treatment/PreT and post-treatment/PostT respectively; see Fig 1B). All the measured variables’ names, and abbreviations if any, are given in Suppl File D1.

### Clustering supports grouping of individuals by disease status and markers by category

The concatenated data of all six cohorts’ data (which are DM*−* Control, DM*−* PreT, DM*−* PostT, DM+ Control, DM+ PreT, and DM+ PostT) was clustered in an unsupervised fashion (i.e., after removing the cohort label to which each individual belongs). This cluster analysis revealed cohort-specific patterns/signatures of markers (see Fig 2) and thereby recovered the grouping of individuals by cohort. That is, almost all individuals clustered based on their disease status, except the DM+ Infected individuals. This cluster analysis was done using data before covariate adjustment (see Methods); repeating this analysis on data after covariate adjustment resulted in a similar trend (see S1 Fig). Furthermore, in Fig 2, we see that the post-treatment samples clustered together with (i.e., exhibited similar marker signatures as) the control samples, especially in the non-diabetic DM*−* backgrounds; this observation is consistent with the effect of the helminth treatment. These observations increased confidence in the quality of the data and allowed us to proceed with further analyses. Clustering of the measured variables (rows of Fig 2) showed that variables that are related to each other by their measurement category are indeed clustered together. For instance, most red blood cell related measurements like Hgb, MCV, MCH, and MCHC (see Suppl File D1 for full names) are grouped closer in the row dendrogram.

**Fig 2.**
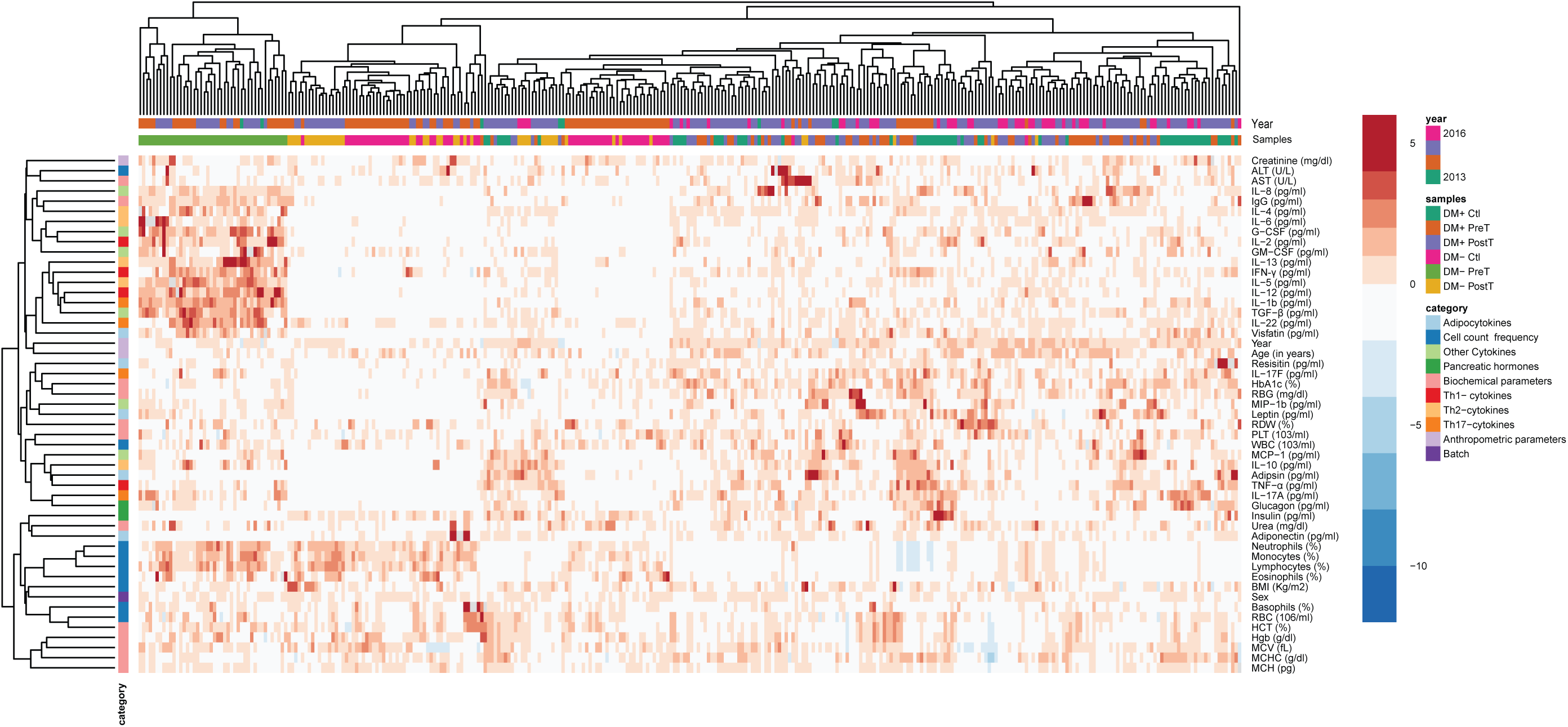
Data overview. Heatmap of the data obtained from concatenating all six cohorts related to helminth/diabetes and performing normalization of each variable (across the samples as explained in Methods; the units of the un-normalized variables are still shown here for reference). Discrete variables like Sex (male or female indicated as 1 or 0 respectively), and Year (when the sample was collected: 2013, 2014, 2015, or 2016) were coded as levels of a factor variable as done in a linear regression model before normalization and visualization.

To better understand the relationship between different variables, besides clustering, we also tried out Bayesian network reconstruction to visualize the measured variables as a network (a set of edges connecting the variables). Network reconstruction was done using data from the before-treatment cohort (see S2 Fig) or the after-treatment cohort (see S3 Fig). In the Bayesian networks, markers that are related by the same measurement category (e.g., most variables in the biochemical parameter category concerning red blood cell and other related measurements like RDW, MCV, Hgb, HCT and PLT) were closer to each other (specifically, lie along a path) in both before and after-treatment cohort Bayesian networks. Also, some of the Th cytokines like IL-17A, IL-2, and IL-1*β* occur together (i.e., lie along a path) in both networks. As a confirmatory check, we inspected the neighbors of the additional disease indicator nodes (called diabetes and helminth status to indicate diabetes and helminth infection respectively). In both the Bayesian networks (S2 and S3 Figs), these disease indicator nodes were indeed connected directly (or via one hop) to the known biomarkers of the disease that were used to define the disease status in our cohorts (biomarker IgG for helminth infection, and biomarkers HbA1c and RBG (random blood glucose) for diabetes [2]). These network-related checks confirmed certain basic expectations about the dataset, and together with the clustering-related observations, encouraged us to proceed to further analyses with confidence. The Bayesian networks can also reveal factors at the interface between the two diseases - for instance, when we looked for a node whose removal would disconnect the helminth and diabetes disease status nodes in the network, we found cytokine GM-CSF in the before-treatment network (and Age in the after-treatment network). While this observation is interesting, we need to view it with caution due to the moderate sample sizes underlying the Bayesian networks. So, a statistical analysis and pipeline that would be valid for our moderate sample size setting is preferable to identify and quantify double-disease markers.

### Identification of DDI markers and their treatment specificity

To identify markers at the interface of helminth infection and diabetes, we applied our DDI pipeline to the helminth-diabetes before- and after-treatment datasets. Our pipeline revealed several DDI markers, each of whose per-disease main and disease-disease interaction effects were significant at the FDR 5% cutoff (see Methods). The results of our DDI analysis are presented as ternary plots in Fig 3, and as statistics corresponding to the main and interaction effects’ tests (p-values and relative proportion of variance of a tested marker explained by single-disease and DDI terms) in Suppl File D2 and Suppl File D3. These results lead to the following findings about the DDI behavior before and after treatment for helminth.

**Fig 3.**
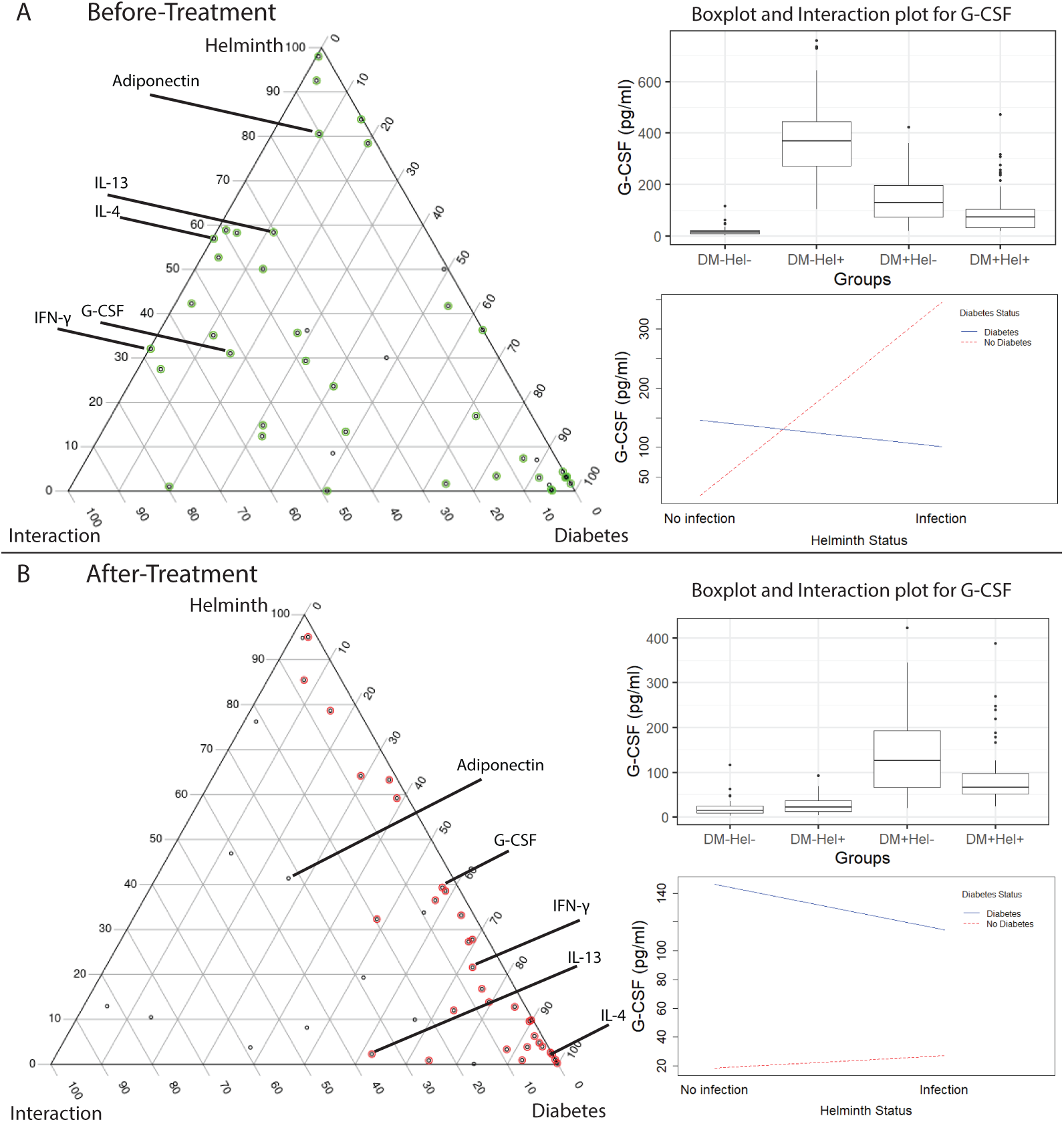
Ternary plots. Ternary plots from diabetes-helminth DDI analysis of the four before-treatment cohorts (A), and the four after-treatment cohorts (B). A black dot (tested variable) with a green/red color around it indicates a main effect variable (at FDR 5%), and a subset of them are further selected as DDI markers (at FDR 5%). Some DDI markers and a non-DDI marker (Adiponectin) in the before-treatment analysis are labeled in both plots. As mentioned in Methods, a ternary plot shows the contribution of the helminth, diabetes, and DDI interaction terms to the explained variance of a marker; for instance, these contributions in plot (A) are respectively 31.04%, 16.46%, and 52.50% for G-CSF (Suppl File D2 and Suppl File D3 show similar percentages for the other variables). (Inset) Boxplot and interaction plot to illustrate G-CSF levels in different cohorts (see S4 and S5 Figs for interaction plots for the other labeled variables in the ternary plot). Interaction plots use lines to show the change in group-specific averages of a variable, with groups defined using helminth and diabetes statuses.

#### Behaviour of DDI markers in before-treatment condition

Our DDI pipeline applied to the before-treatment data revealed 23 DDI markers (at FDR 5%; see Suppl File D2), with IL-17A, Visfatin, and IFN-*γ* being the top 3 DDI markers based on the relative contribution of the interaction (DDI) term to explaining the individual-to-individual variation in these cytokines (also referred to simply as the term contribution hereafter; see Methods). The identified DDI markers (see Suppl File D2) included certain DDI markers like IL-4 and IL-8 that are not affected by diabetes in isolation (diabetes term contribution of less than 5%), but are highly influenced by diabetes during helminth comorbidity (DDI interaction term contribution of more than 40%); these cytokines also have a high helminth term contribution of more than 50% consistent with earlier studies (e.g., IL-4 discussed in [6]). Viewing an interaction conversely, we found several Th2 cytokines known to be highly expressed during helminth infection such as IL-4, IL-5, and IL-10 to also be influenced by concomitant diabetes (i.e., they had high DDI interaction term contributions of 42.79%, 36.51%, and 55.52% respectively).

Since inflammation-related cytokines associated with diabetes are known to be downregulated during helminth infection, we checked their DDI statuses and indeed found the DDI term contributions of inflammatory cytokines IFN-*γ* (see Fig 3A), TNF-*α*, IL-2, and IL-17A to be high (67.68%, 48.88%, 54.13%, and 79.57% respectively). This downregulation could be due to the opposite effect of Th2 cytokines associated with helminth response on the pro-inflammatory Th1 cytokines (IFN-*γ*, TNF-*α*, and IL-2) and Th17 cytokine IL-17A [17, 18]. Taken together, these results on the relative contribution of single- and double-disease influences on different types of DDI markers offer a data-driven quantitative view of helminth-diabetes comorbidity.

#### Majority of the DDI markers resolve in after-treatment condition

Our pipeline applied on the after-treatment data revealed 8 DDI markers at FDR 5% (Suppl File D3 and Fig 3B). Compared to the 23 DDI markers discovered under the before-treatment condition, this shows that the DDI effect of many variables was lost due to the anthelmintic treatment. While this is to be expected, our analysis provides a quantitative estimate of the DDI loss (Suppl File D3 Table, compared with Suppl File D2). For instance, focusing on the same sets of cytokines discussed above, the respective DDI term contribution of Th2 cytokines IL-4, IL-5, and IL-10 decreased to 0.05%, 0.19%, and 8.46%; and that of Th1 cytokines IFN-*γ*, TNF-*α*, and Th17 cytokine IL-17A decreased to 6.09%, 14.56% and 5.96% respectively in the after-treatment condition. These results offer a quantitative view of the observations made in our earlier study [2]; and is also consistent with the treatment-driven decrease in the levels of Th2 cytokines [13, 14].

### Replication of DDI signals in independent cohorts

Having identified DDI markers in the four-cohort (before-treatment) discovery dataset, we would like to test if the interaction patterns replicated in a validation dataset obtained from different cohorts. The validation dataset had all but one of the four cohorts required to do DDI analysis (the Hel+DM*−* group was missing); so we had to improvise and test for replication only the DDI signatures that are visible in the three available validation cohorts. First, we note that a majority of the DDI makers identifed in the discovery dataset and discussed in the text above (specifically the Th1, Th2, Th17, and top-3 DDI term contribution cytokines) show largely similar expression patterns between the discovery vs. validation datasets on a visual inspection (see Fig 4A and S6 Fig).

**Fig 4.**
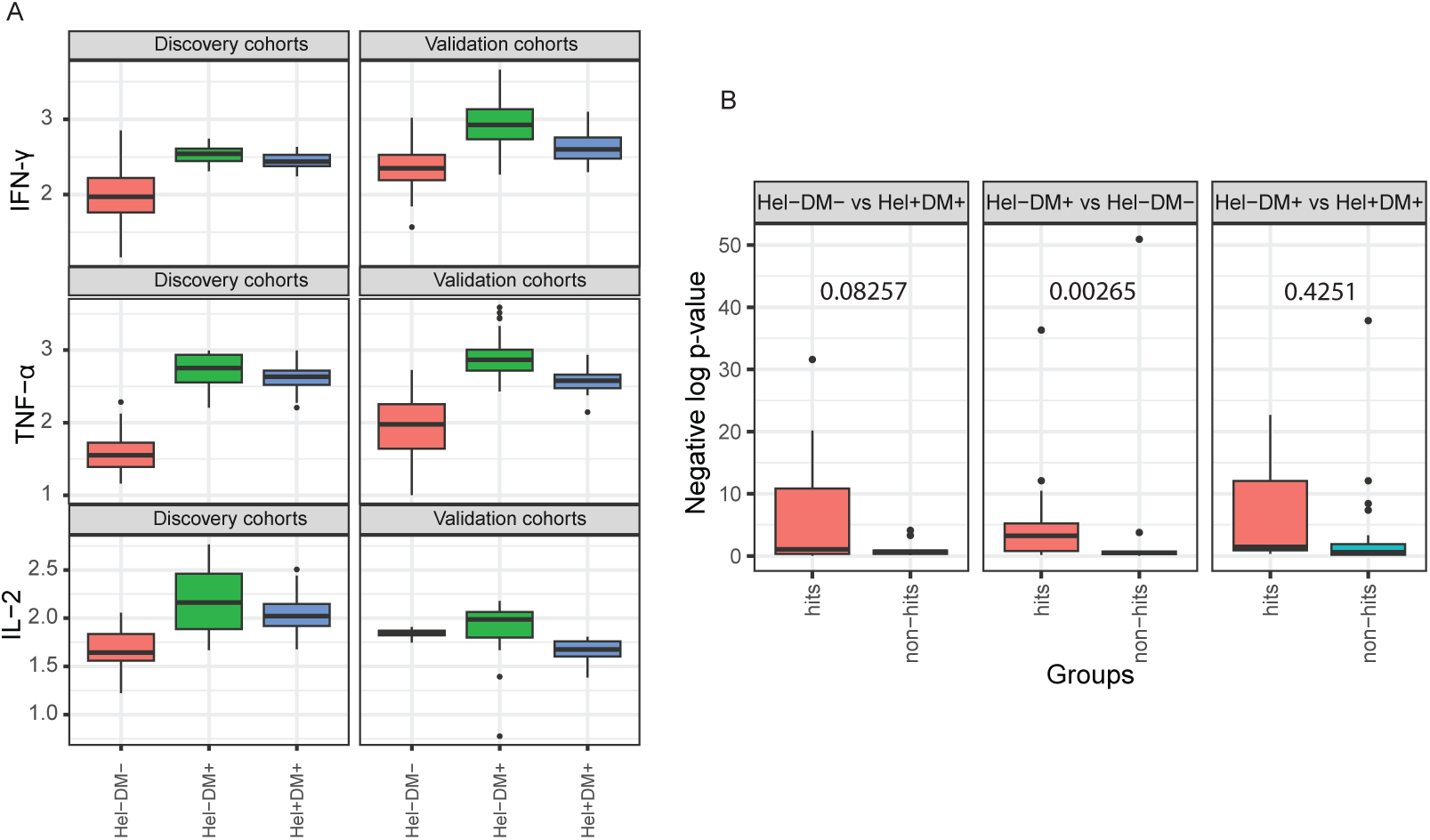
Replication of DDI patterns and DE signals. (A) Expression levels of certain Th1 cytokines are shown after (base-10) log transformation in both discovery and validation cohorts. See S7 Fig for similar boxplots for other (Th2, Th17, etc.,) cytokines. (B) DE signals (*−log*_10_(DE p-value)) in validation dataset of the DE hits vs. non-hits that were identified in the discovery dataset. Whether DE signals of hits are better than that of non-hits was tested using a robust t-test and its p-value is shown for each comparison.

To quantify this visual inspection of replication of expression patterns of DDI markers, we performed three differential expression (DE) analyses comparing two of the three available cohorts at a time (see Methods). Each DE analysis yielded hits that are DE markers called at a specific adjusted DE p-value threshold of 0.0001 in the discovery dataset, and non-hits that are the remaining tested variables (see Methods). We can now inspect whether hits have higher DE signal in the validation dataset compared to non-hits, and we do find that to be the case for two of the three DE analyses (according to robust t-test p-value cutoff of 0.1; see Fig 4B). This trend for hits defined using an adjusted DE p-value cutoff of 0.0001 in the discovery dataset also held when this cutoff was changed to 0.01 and 0.05 (see S8 Fig; Suppl File D4 lists the hits and non-hits for the different cutoffs used). These results taken together support that the cohort-specific expression and DE-related patterns of DDI/other markers in the discovery dataset do replicate in the validation dataset.

### Interpreting celltype-specific DDI markers

Most of our DDI markers are cytokines that play the role of mediators between different immune celltypes. We would like to query external databases on cytokine-celltype and biological pathways to hypothesize which immune celltypes and biological processes in these celltypes are associated with the DDI cytokines. Note that a particular cytokine may be produced/secreted by a source celltype to elicit a response from other target immune cells (of the same or different celltype).

To determine the likely source cells of DDI cytokines, we query the corresponding DDI genes in a gene signature matrix called LM22, which contains the expression of 547 genes across 22 immune celltypes [19]. Fig 5 shows the expression patterns of seven of our DDI genes that overlap with the 547 LM22 genes. High expression of DDI marker genes GM-CSF and IL-5 in memory-activated CD4 and mast cells suggests that these may be the source immune celltypes of these cytokines. We can observe the same for pro-inflammatory cytokines like IFN-*γ* and TNF-*α*. In contrast, the expression of these cytokines in CD4 naive cells is low.

**Fig 5.**
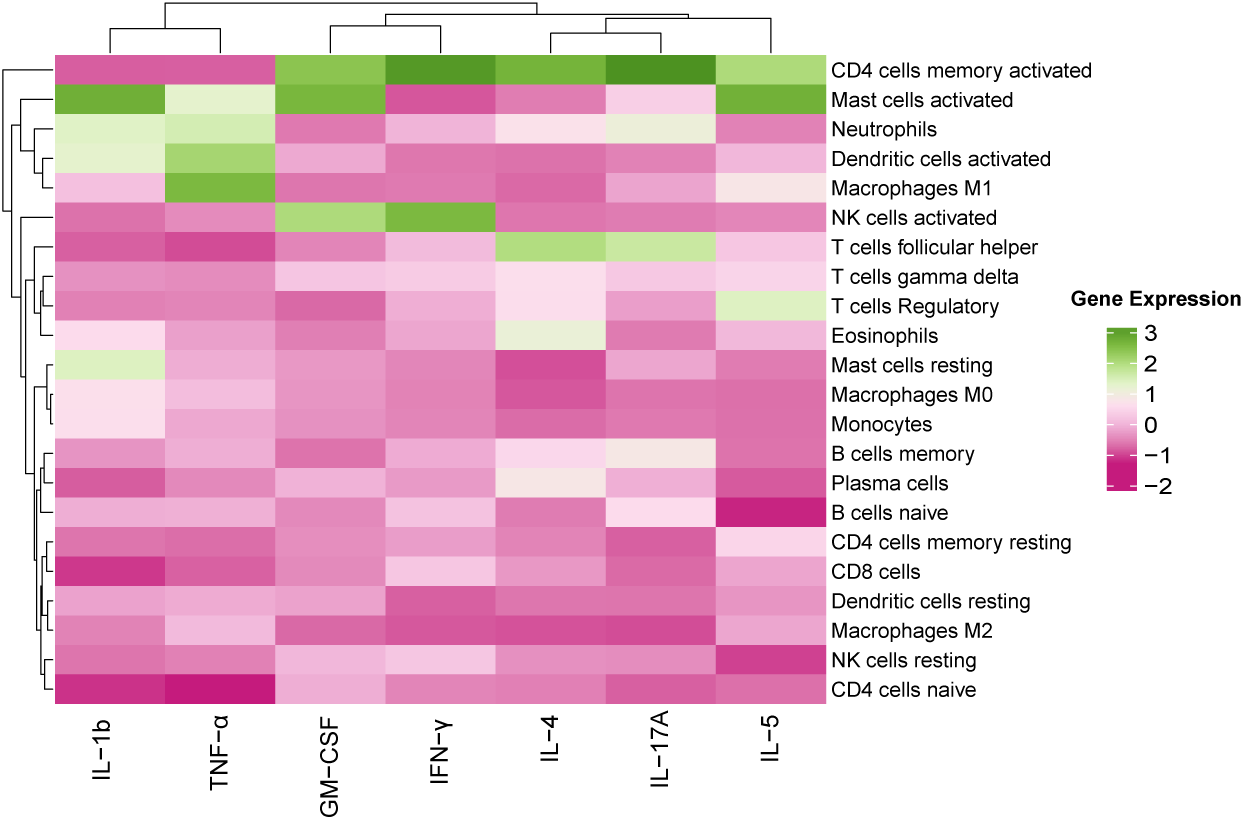
Expression of DDI markers in different immune cells. The cell type specific expression patterns of DDI genes (encoding the DDI cytokines identified using our proposed pipeline) that overlap with the 547 genes in the signature matrix LM22 are shown. The heatmap is column-normalized, hence the green shades represent higher expression levels compared to pink shades in each column.

Having predicted the likely source cells of DDI markers, we would like to next predict their effect on target cells, focusing on the genes and pathways targeted by these DDI cytokines. Towards this, we employ web-based tools CytoSig and WebGestalt to identify which biological pathways (Reactome pathways) are enriched for genes targeted by each DDI marker of interest (specifically focusing only on the Th-related DDI cytokines; see Fig 6). Two of the key findings from this analysis follows. First, pathways mediated by Th2 cytokines like IL-10 Signaling and IL-4/IL-13 Signaling were enriched for genes affected by the DDI markers. Further, IL-4/IL-13 cytokines are essential for the proliferation of immune cells like macrophages [20, 21] (see Discussion for details). Second, pathways related to GPCR ligand binding and Class A/1 (Rhodopsin-like receptors) were also enriched for genes responding to the DDI markers IL-1, TNF-*α*, and GM-CSF. These receptors are responsible for cell differentiation and chemotaxis of immune cells like monocytes, neutrophils, and macrophages [22]. Therefore, these findings on DDI Th cytokines reveal not only their target pathways, but also their (above-mentioned) target immune celltypes.

**Fig 6.**
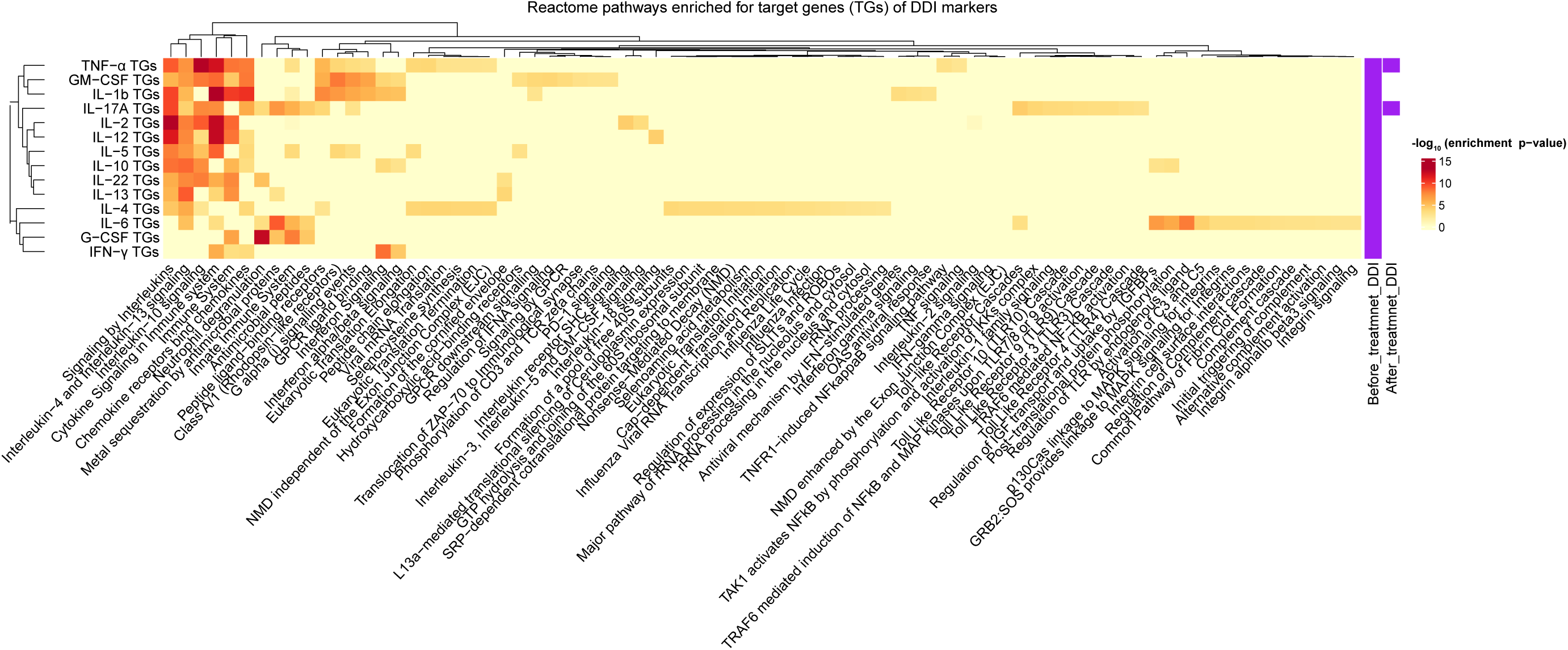
Pathways targeted by DDI markers. Pathways (columns) that are enriched for target genes (TGs) of DDI markers (rows, focusing on DDI Th cytokines). Extra two columns at the end indicate the DDI markers in before- and after-treatment data. The heatmap shows the negative of the base-10 logarithm of the (multiple-testing) adjusted enrichment p-value (i.e., p-value of enrichment of the Reactome pathway for the DDI cytokine’s TGs; see Methods for details). NMD, Nonsense Mediated Decay; IGF, Insulin-like Growth Factor; IGFBP, Insulin-like Growth Factor Binding Proteins.

## Discussion and Conclusion

### Summary of our findings

In this work, we propose a three-step pipeline for understanding the interface between any two diseases for which multi-cohort measurements are available. We apply this pipeline to perform a systematic quantitative analysis of the interaction between diabetes and helminth. We identified DDI markers through main and interaction analysis and quantitated the relative contribution of the per-disease statuses (helminth and diabetes terms) and disease-disease interaction (DDI) terms to explain the individual-to-individual variation of the measured variables (adipocytokines, Th1, Th2, Th17 cytokines, other cytokines, pancreatic hormones, biochemical parameters, cell count frequencies, and anthropometric parameters). The identified DDI markers such as IFN-*γ*, TNF-*α*, and IL-2 at the interface of both diseases show similar signatures of interaction in multiple cohorts of discovery and validation data, thereby increasing confidence in our findings. We also interpret certain DDI cytokine markers in terms of the source immune celltypes that likely secrete them and target pathways like IL-10 and IL-13/4 signaling that likely respond to these cytokines.

There have been human and animal studies that have analyzed diabetes and helminth infection separately [23, 24], and also a few that have analyzed the co-occurrence of these two diseases [2, 25] – our study’s key contribution is in providing a quantitative framework/pipeline for identifying and interpreting DDI markers. To elaborate, cross-sectional epidemiological studies and experiments in animal models support the potential protective effects of helminth infections on the development of type-2 diabetes. These beneficial effects seem to stem from the helminth-induced downregulation of inflammatory pathways involved in the development of insulin resistance or diabetes [2, 26]. This double-disease downregulation is largely consistent with the DDI term contribution percentages that we estimate for inflammatory cytokines like IFN-*γ* and TNF-*α* (Suppl File D2). Furthermore, our pipeline also revealed other DDI markers like IL-6 and IL-10, which are known risk factors for diabetes [27]; and TNF-*α*, IL-12, IL-2, and G-CSF, which are shown to have physiological effects on diabetic individuals [28].

Besides quantifying double-disease interaction effects, our study also quantifies per-disease effects such as the helminth infection’s relative contribution to the plasma levels of different markers. The high-to-moderate helminth term contribution (expressed as percentages in Suppl File D2) of certain biomarkers related to Th2 responses, like IL-4 (56.94%), IL-5 (50.08%) and IL-10 (12.40%), are consistent with earlier observations of elevated plasma levels of these markers during helminth infection [12]. The drop in helminth term contributions of IL-4, IL-5, and IL-10 to less than 5% in after-treatment data (Suppl File D3) is also concordant with our earlier observations on the downregulation of these markers after anthelmintic treatment of helminth-infected individuals [25].

Regarding the effect of anthelmintic treatment, we provide a comprehensive across-all-markers view of the shift (decrease) in their interaction effects from the before- to the after-treatment ternary plots. For instance, the DDI term contribution of pro-inflammatory markers like IFN-*γ*, TNF-*α*, and IL-17A decrease after treatment. We observed that these cytokines show a marginal increase in expression in the post-treatment individuals, but they do not revert to their original control levels. The reasons for this memory of helminth infection in the body would be interesting to explore in the future.

### Additional biological information on the DDI markers

Our Reactome pathway enrichment analysis revealed IL-10 and IL-4/13 signaling pathways to be targeted by multiple DDI cytokines. We now provide literature support to clarify the activation of these signaling pathways in immune cells like macrophages under helminth infection. Turner et al. have demonstrated in mouse models that exposure to IL-4/IL-13 cytokines fosters the proliferation of macrophages (M0) to alternatively-activated macrophages (AAM/M2) [20]. AAM/M2 macrophages, which differentiate from monocytes and macrophages upon activation of IL-4/13 signaling pathways, are required to fight against helminth infection in the target tissue [21]. In their work, Berbudi et al. suggest a potential mechanism for how IL-4/13 signaling pathways activate AAM/M2 macrophages [15]. These AAM/M2 macrophages are also known to produce high levels of IL-10 [29, 30], which in turn can activate IL-10 signaling in macrophages. Hence macrophages can serve both as the source immune celltype for IL-10 and as the target celltype where IL-4/13 signaling and subsequently IL-10 signaling are activated when exposed to the relevant DDI cytokines.

### Caveats, Merits and Future directions

There are some caveats to our analysis. While we’ve adjusted our data for known confounding factors to mitigate bias in our identified DDI markers, unknown confounding factors cannot be ruled out; so our DDI methodology should be viewed as generating hypotheses (DDI markers) that need to be further experimentally validated. Our current study focuses on each variable separately to test if it is a DDI marker to simplify analysis and interpretation. However many markers are correlated and therefore a per-marker analysis cannot capture correlated trends. In the future, we could build a combined model that takes multiple markers as input to predict the disease status as output and use the feature importance scores derived from this model to quantify the aggregate DDI effect of a set/module of measured variables. Another caveat concerns the replication testing of observations from our four-cohort discovery dataset using only a three-cohort validation dataset. Although we could not perform an identical interaction analysis on the validation dataset (due to the non-availability of one of the four cohorts in the validation dataset), the aggregate interaction signatures (based on pairwise differential expression analyses of hits) in the discovery dataset did replicate in the validation dataset. One more limitation of the interpretation step in our DDI pipeline is the inability to predict rare cell types as sources of the DDI cytokines. We did not focus on rare cell types, because we wanted to employ an established signature matrix called LM22 from a popular method CIBERSORT to predict the source cells. Whereas LM22 contains the gene expression of 22 major immune cell types only, if other signature matrices involving rare cell types become available and established in the future, we can also expand our source predictions to rare cells in the future.

One of the merits of our work is that provided there are four sets of cohorts (two single-disease, double-disease, and control cohorts), our DDI pipeline and associated analyses could be carried out to understand biochemical variables at the interface of any pair of two diseases of interest. The results from the Helminth-Diabetes interaction show a concrete application of our DDI methodology to uncover cytokine and related markers that play a role in the interface of these two diseases. The DDI markers are the molecules (in our case, mostly cytokines) that are affected in a different way (synergistic or antagonistic way) in the comorbid condition relative to the individual disease conditions. As such, the DDI markers and the biological pathways that they are linked to could be targeted for novel comorbid disease therapy. Given that the DDI markers are found through associations, it is possible that they cause the disease or are a response to it. In the former case, the novel treatment could focus on curing the causes of the diseases, whereas in the latter, the treatment could focus on alleviating the symptoms of the comorbid condition.

In the future, our DDI methodology could be applied to analyze any two diseases and any set of variables (such as flow cytometry-based measurements of cell surface markers or gene expression levels in transcriptomics datasets). For instance, it could be applied to the multi-cohort gene expression data available for the disease pair, diabetes and pancreatic cancer [31]; gene expression and blood-based measurement data available for diabetes and tuberculosis [32], or metagenomic and the associated species abundance data available for samples affected with combinations of two parasites [33]. Application of our pipeline on these datasets will help to understand the markers at the interface of the two co-occurring diseases, and suggest targets for new treatment strategies that address double-disease conditions differently than simply prescribing treatments developed separately for each disease.

## Materials and Methods

### Our DDI Methodology/Pipeline (DDI-P)

As indicated in Fig 1A, our DDI pipeline comprises three different steps explained in detail in this section. Briefly, the first step is preprocessing and exploratory visualization/analysis of our multi-cohort data pertaining to two diseases. After that, the DDI markers are identified using a main and interaction effect analysis in the second step; and interpreted using prior biological knowledge (such as stored in cytokine knowledge bases) in the final third step of the pipeline.

To ensure that the detected DDI markers are relevant for the two diseases under consideration, we recommend that the four groups (control, disease 1 only, disease 2 only, and comorbid group) be matched on possible confounding factors and covariates. For instance, it is preferable that all four groups have similar age distribution, sample sizes, etc., and that the three disease groups have similar disease severity levels. However, it may be hard to get access to such cohorts, and any differences in the sample size or covariate distributions can be corrected/adjusted in the data using linear regression models.

#### DDI-P (i): Dataset preprocessing and visualization

Given data from control, single-disease (disease 1 and disease 2, such as diabetes and helminth infection), and double-disease cohorts, this step concatenates data on the same set of demographic/biochemical variables measured across these four cohorts. We also add disease status variables D1 and D2. For an individual, D1 is a binary indicator variable of disease 1, i.e., we set D1=1 if the individual has disease 1, and D1=0 otherwise. D2 is similarly a binary indicator variable of disease 2. For individuals who suffer from both diseases, their D1=1 and D2=1. The concatenated data from the four cohorts can be normalized later depending on the analysis done. For the cohorts mentioned in this paper, normalization (standardization, i.e., centering and scaling) for all the variables across the cohorts was carried out only to plot the heatmap. Heatmap visualization of the concatenated normalized data can reveal the expression patterns of all variables across the four cohorts, and help us understand the data-driven grouping of individuals (and variables) in the dataset into clusters with similar patterns.

#### DDI-P (ii): Main and interaction effect analysis

To gain better insight into markers at the double-disease interface and to specifically identify DDI markers, we performed main effect (per-disease contributions) analysis, followed by interaction effect (double-disease contributions) analysis on the concatenated data from the four cohorts.

##### Linear regression models

In the main effect analysis, a linear regression model of the dependent variable (e.g., cytokine) is fit using only the independent variables/features: D1, D2, and all covariates. Whereas in the interaction effect analysis, a linear regression model of the dependent variable is learnt by including an additional interaction variable/feature, which captures the combined effect of D1 and D2 (coded as an interaction term D1:D2, which is set to D1*D2, the product of D1 and D2) [34, 35]. These linear regression models and a baseline control model (shown below) are fit to the data, and compared using statistical tests to find the main and interaction effect p-values, and subsequently the DDI markers.

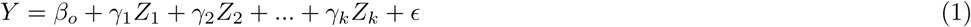

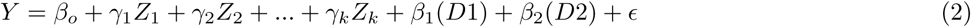

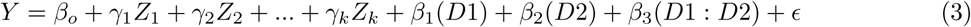

Here, *Z*_1_,*Z*_2_,..,*Z_K_* are the K covariate variables measured in the cohorts and *γ*_1_,*γ*_2_,..,*γ_K_* their corresponding coefficients. *Y* is the dependent variable, and the terms D1, D2, and D1:D2 discussed above are independent variables. The unexplained variance in these linear regression models is modeled by Gaussian noise terms *ɛ*. Here *β*_0_ is the intercept term, *β*_1_, and *β*_2_ are the weights associated with the per-disease terms and *β*_3_ is the weight associated with the double-disease interaction term.

The rationale for choosing the above linear regression models with main and interaction effects is based on two primary considerations: sample size requirements and ease of interpretation. The coefficients (effect sizes) in linear models can be reliably estimated from moderate sample sizes, and these estimated coefficients are also easier to interpret due to additive contributions of the different terms. Furthermore, non-additive effects (e.g., sum of individual effects of two diseases differing from the joint comorbid effect of both diseases) can be naturally modeled using interaction terms in linear models.

##### Statistical tests to identify a DDI marker

The main effect analysis was performed by comparing the two nested linear models, Eq 2 vs. Eq 1 above, using a F-statistics based test (implemented in anova.lm function in R); and adjusting/correcting the resulting p-values for multiple testing using the Benjamini-Hochberg FDR procedure (implemented in p.adjust function in R). Using an adjusted p-value cutoff of less than 0.05 (i.e., at FDR 5%), the variables that show the main effect were selected. The same procedure was repeated on the selected main effect variables alone to perform interaction analysis in order to obtain the interaction effect p-values (also known as DDI p-values) and variables, but here the compared nested linear models are Eq 3 vs. Eq 2 instead (and multiple testing correction is done only over the tested main effect variables). A marker that gets selected as main effect and interaction effect variable in the statistical tests above, each at FDR 5%, is called a disease-disease interaction (DDI) marker. Note that this two-step strategy, viz., identifying variables that exhibit a main effect followed by testing only the main effect variables for interaction effects, is a standard way to reduce multiple testing burden when detecting interaction effects.

##### Ternary plot

The results from interaction analysis are visually represented through a ternary plot [5, 36]. Each vertex in the ternary plot represents the three terms: D1 (disease 1), D2 (disease 2), and D1:D2 (interaction or DDI), and each dot in the plot a tested marker. For a tested marker, the relative proportion of the marker’s variance explained by these three terms are used as coordinates to mark the variables in the ternary plot. The black dots in a ternary plot are all variables considered/tested in the study, and a green/red circle around a black dot indicates that the variable is also a main effect variable. TernaryPlot() function from the package Ternary was used to generate the plots in Fig 3.

#### DDI-P (iii): Interpretation of DDI markers

We would like to find out biological pathways that are targeted by a DDI cytokine of interest, i.e., pathways in a cell that respond when the cell is exposed to that cytokine. To predict such pathways, we first collect known target genes (TGs) of the cytokine from the CytoSig database [37] and then test different pathways for enrichment of these query TGs using the WebGestalt tool [38]. In a bit more detail, for a DDI cytokine of interest (such as a Th-cytokine DDI marker), we used CytoSig to retrieve the top 100 entries (search results) of TGs for the cytokine (see Suppl File D5 for TGs of some DDI cytokines) and used WebGestalt ORA (Over-Representation Analysis) to test enrichment of this query set of TGs in each pathway from the Reactome database. WebGestalt function from the R Package WebGestaltR (version 0.4.6) was employed here, with its key function arguments set as follows: “pathway REACTOME” for enrichDatabase, “genome” as referenceSet, and “genesymbol” as interestGeneType. WebGestalt can adjust the enrichment p-values of a cytokine for multiple testing (across all tested Reactome pathways), and apply a cutoff of 0.05 on the adjusted p-values to obtain the Reactome pathways significantly enriched for the cytokine’s TGs at FDR 5%; these pathways enriched at FDR 5% are referred to as the target pathways of the tested cytokine.

To visualize the above pathway enrichment results as a heatmap, we take the union of target pathways of all tested cytokines, and retrieve their enrichment p-values to assemble an enrichment signal matrix. This matrix was input to Heatmap from ComplexHeatmap (version 2.14.0) to obtain the heatmap in Fig 6. In detail, the (*i, j*)-th entry of the enrichment signal matrix is simply the negative log (to base 10) of the multiple-testing-adjusted enrichment p-value of pathway *j* for the top TGs of cytokine *i* (see Suppl File D6). To assemble this matrix, we need to retrieve the enrichment p-values of not just the significantly enriched pathways of a tested cytokine, but also other pathways; so we reran the same WebGestalt function above, but this time, not with a FDR cutoff of 0.05, but with the following arguments: 1 as minNum, 2000 as maxNum, 10000 as topThr, “top” as sigMethod, and 40 as reportNum. We then corrected the resulting p-values for multiple testing using the Benjamini-Hochberg method (implemented using the p.adjust function in R) to get the adjusted enrichment p-value.

To determine the likely source cells of the DDI cytokines (specifically those DDI cytokines that were part of the CytoSig/WebGestalt-based enrichment analysis above), we query the corresponding DDI genes encoding these cytokines in a gene signature matrix called LM22. The LM22 matrix containing the expression of 547 genes across 22 immune celltypes [19] was retrieved from the R package ADAPTS (version 1.0.22). However, only a subset of the queried DDI genes may overlap with the 547 LM22 genes. The submatrix of LM22 over these overlapping genes is input to the Heatmap function from the ComplexHeatmap package (version 2.14.0) to obtain the desired heatmap (such as the one in Fig 5).

### Application of DDI-P to Helminth-Diabetes cohorts

We applied DDI-P to perform interaction analysis of helminth infection and diabetes disease, with the cohorts pertaining to different combinations of these two diseases (see Fig 1B). These cohorts were studied and profiled at the National Institute for Research in Tuberculosis (NIRT) located in Chennai in South India.

#### Helminth-Diabetes cohorts: Dataset, covariates, preprocessing and visualization

##### Dataset overview

All the samples used in this study with Type 2 Diabetes Mellitus (Diabetes) and with/without Strongyloides stercoralis (Ss Helminth) infection were profiled to make blood-based measurements and the resulting data analyzed to detect disease-specific expression patterns in our earlier work [2]. We provide a comprehensive and quantitative DDI pipeline based interaction analysis of all the profiled marker variables in this data in the current work. This data was collected from 118 individuals consisting of 58 individuals with no helminth infection (DM+ Control) and 60 helminth-infected individuals (DM+ Pre-treatment). Data analyzed here additionally include measurements from individuals without diabetes and with/without helminth infection collected following the same procedure as detailed in our previous work [2]. Both datasets contain measurements of the same set of demographic variables, biochemical parameters, cytokines, hormones, etc. (with all the measured variables used in this study, along with their category, listed in Suppl File D1). As mentioned before in Fig 1B and in the Results section, data without diabetes was also collected from 120 individuals, including 60 individuals with no helminth infection (DM*−* Control) and 60 helminth-infected individuals (DM*−* Pre-treatment). All helminth-infected individuals were subjected to anthelmintic treatment, and measurements were repeated six months later (DM+ Post-treatment, DM*−* Post-treatment); with the exception of a subset of individuals who couldn’t be followed up for measurements in DM*−* Post-treatment. Note that DM*−* Post-treatment contains 44 individuals, and so the DM*−*Pre-treatment data (containing 60 individuals) was restricted to the same 44 individuals when doing all DDI and other covariate-related analyses in this paper.

##### Covariates selected

A set of confounding factors or covariates that applies across our helminth-disease cohorts need to be selected, before we can proceed with all the DDI or related downstream analyses in our DDI pipeline. For this reason, we chose certain demographic characteristics like Age (in years), Sex (male/female), and BMI (Body Mass Index in Kg/*m*^2^), technical factor like Year (Year in this paper refers to the year of sample collection: 2013, 2014, 2015, or 2016), and physiological state indicators like Kidney function indicator (Creatinine (mg/dl)), and Liver function indicators (Aspartate aminotransferase (AST in (U/L) and Alanine aminotransferase (ALT in (U/L)).

For the different cohorts analyzed in this study, these seven covariates (Age, Sex, BMI, Year, Creatinine, AST and ALT) were distributed in a largely similar fashion, but with some notable differences (S1 Table) which could be adjusted for prior to doing DDI or downstream analyses. This adjustment will make it more likely that any differences between the cohorts is due to differences in the status of the two diseases in these cohorts, rather than differences in demographic or other such factors between the compared cohorts.

##### Clustering-based visualization of dataset

The function Heatmap from ComplexHeatmap was used to generate heatmaps of the concatenated data from the six helminth-diabetes cohorts mentioned above. Samples were clustered according to the Optimal Leaf Ordering method. The results in Fig 2 were generated using the raw data without any adjustment for the covariates (but with marker variables being normalized, specifically standardized, across the samples). Covariate-adjusted data is simply the residuals obtained from a linear regression model given by Eq 1 (i.e., using the marker expression as the dependent variable and the seven covariates discussed above as independent variables) fitted to concatenated data from all six cohorts; this covariate-adjusted data was standardized before plotting as a heatmap in S1 Fig.

##### Bayesian Network (BN) reconstruction from dataset

BN reconstruction was done to supplement the clustering-based exploratory analysis/visualization. Two BNs were reconstructed, one using the before-treatment cohorts and another using the after-treatment cohorts. BN or simply Network here refers to the set of edges connecting the nodes (measured variables) in the network and can be reconstructed or learnt from data based on the (marginal/conditional) correlations between different variables in the data. Here, the helminth and diabetes statuses (D1 and D2) are discrete variables, while the remaining variables are continuous variables. R package bnlearn (version 4.6) was used to learn the directed acyclic graph (DAG) structure of the networks from the above-mentioned datasets [39]. We used boot.strength() to estimate the strength of the learnt arcs in the networks. In this process, we generate several bootstrap resamples of the data, and learn separate DAG-structured networks from each such bootstrap-resampled dataset (using the Min-Max Hill Climbing (MMHC) hybrid structure learning algorithm). Across all bootstrap-resampling-based networks, we can now count how often each arc gets repeated and express it as a fraction from 0 to 1, and then obtain a consensus network with arcs that appear often using averaged.network() (all of these functions were run with the default input arguments). The final BN is the averaged consensus network (dropping direction of arcs in the graph since it is a consensus network) containing only arcs whose strength is above a particular threshold value (0.5). The BN reconstructed from before-treatment data is in S2 Fig (and after-treatment data is in S3 Fig).

#### Helminth-Diabetes cohorts: Interaction analysis

To have a better insight into the variables involved in the double-disease interface both before and after treatment for helminth, main and interaction analyses were performed on the before-treatment and the after-treatment cohorts. Towards this, DM*−* Control, DM*−* Pre-treatment, DM+ Control, and DM+ Pre-treatment samples were considered for the before-treatment interaction analysis. DM*−* Control, DM*−* Post-treatment, DM+ Control, and DM+ Post-treatment samples were considered for the after-treatment analysis.

The main/interaction effect analysis mentioned in the DDI pipeline was applied once on the before-treatment data and another time on the after-treatment data for each tested marker. The same seven covariates used in cluster analysis of covariate-adjusted data mentioned above (i.e., Age, Sex, Year, BMI, Creatinine, AST, and ALT) was also used as covariates in the DDI pipeline analysis. Suppl File D2 lists the parameters of the linear models estimated during the main and interaction effect analyses of the before-treatment data (and Suppl File D3 presents similar estimates for the after-treatment data). The DDI (or other) term contributions of the before- vs. after-treatment data (Suppl File D2 vs. Suppl File D3) can then be compared to understand the effect of helminth treatment on the DDI status of markers. Note that the DDI term contribution of a marker is the relative contribution of the DDI term to the *explained* variance of the marker; still it is valid to compare these relative contributions after and before treatment for the following reason. For a majority of the markers, the fraction of the marker’s variance that is *unexplained* is not very different between after- and before-treatment data (S6 Fig).

#### Helminth-Diabetes cohorts: Interpretation of DDI markers

The downstream analyses (target pathway enrichment analysis using CytoSig/WebGestalt and source celltype prediction using LM22 matrix) for interpreting DDI markers, was also performed on the identified helminth-diabetes DDI markers. Only the DDI markers G-CSF, GM-CSF, IL-1*β*, IL-2, IL-4, IL-5, IL-6, IL-10, IL-12, IL-13, IL-17A, IL-22, TNF-*α*, and IFN-*γ* (rows of Fig 6) were considered for the target pathway enrichment analysis, since CytoSig did not have any information on the rest of the helminth-diabetes DDI markers. Further, only 8 of these markers (columns of Fig 5) overlapped with the genes in the LM22 matrix, and hence the source celltype analysis was also performed only for these 8 helminth-diabetes DDI markers.

### Replication analysis

#### Validation cohorts overview

The replication analysis was carried out using validation cohorts pertaining to a different study at NIRT. The validation dataset consisted of measurements from 31 non-diabetic, non-helminth-infected individuals as controls (denoted DM*−* Control, or DM*−*Hel*−*), 43 diabetic non-helminth-infected individuals (denoted DM+ Control, or DM+Hel*−*), and 41 diabetic helminth-infected individuals (denoted DM+ Pre-treatment, or DM+Hel+). The validation study had a different design, and so didn’t have non-diabetic helminth-infected individuals and also didn’t have post-helminth-treatment data. Of the 50 markers measured in the discovery cohorts, 42 were also profiled in the validation cohorts, and we focused on these common markers for conducting the replication tests.

#### Replication testing based on DE analyses

Interaction analysis to discover DDI markers requires data from four groups of individuals (the two single-disease, one double-disease, and one control groups). If a validation study has fewer groups, like three instead of four in our case as described above, we can still validate/verify certain interaction signatures of the markers as follows. We performed 3 different pairwise comparisons of the 3 groups as described below, using the discovery dataset first and the validation dataset next.

1. DM*−* Control vs. DM+ Pre-treatment (Hel*−*DM*−* vs. Hel+DM+). This comparison captures double-disease effects relative to controls.
2. DM+ Control vs. DM+ Pre-treatment (Hel*−*DM+ vs. Hel+DM+). This comparison captures helminth effect under a diabetes background.
3. DM*−* Control vs. DM+ Control (Hel*−*DM*−* vs. Hel*−*DM+). This comparison captures diabetes effect under a non-helminth background.

We conduct DE analysis on each pairwise comparison above using the discovery cohorts first. The DE analysis is essentially a main effect analysis where the dependent variable (marker expression) is regressed over the covariates, diabetes term, and helminth term, and the main effect or DE p-values calculated using a F-statistic based test (cf. Eq 2 vs. Eq 1 statistical test discussed above). The p-values were adjusted for multiple testing across all markers using the Benjamini-Hochberg procedure. Hits are markers/variables whose adjusted DE p-value is at most a stringent cutoff of 0.0001; and non-hits are the rest of the tested markers (other relaxed cutoffs of 0.01 and 0.05 were also tried and resulted in similar trends).

Having identified the hits and non-hits using the discovery cohorts’ data, we would now like to test it for replication in the validation cohorts. Towards this, we inspect the DE signal (*−log*_10_(DE p-value)) of hits vs. non-hits in pairwise comparisons of the validation cohorts (see Fig 4). If this DE signal of the hits are better than that of non-hits (as tested using Yuen’s robust t-test, implemented in the yuen function from the WRS2 package in R [40]), we consider the DE signals of hits detected in the discovery dataset as replicated in the validation dataset. Yuen’s variant of the t-test was chosen due to its robustness against outliers.

## Declaration

### Ethics approval and consent to participate

All participants were examined as part of a natural history study protocol (12-I-073) approved by Institutional Review Boards of the National Institute of Allergy and Infectious Diseases (USA) and the National Institute for Research in Tuberculosis (India), and informed written consent was obtained from all participants.

### Availability of data and materials

We have publicly released the code implementing our DDI pipeline, and the code to reproduce the figures/tables and supplementary figures/tables in the manuscript at https://github.com/BIRDSgroup/DDI. The data pertaining to the six helminth-diabetes cohorts are available from the authors upon request.

### Authors’ contributions

NS and MKN conceived and developed the DDI methodology; NS implemented the methodology and applied it to diabetes-helminth multi-cohort data to obtain results; NS and PP preprocessed and performed exploratory analysis of the multi-cohort data; NS, AR, NPK, SB,and MKN interpreted the results; AR, NPK and SB generated the multi-cohort data, with SB leading and supervising this experimental effort; NS and MKN wrote the manuscript, and all authors reviewed the manuscript; MKN led and supervised the overall project.

## Acknowledgements

We thank members of our BIRDS (Bioinformatics and Integrative Data Science) research group, and members of IBSE (Center for Integrative Biology and Systems Medicine) for their valuable input during the presentations of this work. The research presented in this work was supported by the Wellcome Trust/DBT grant IA/I/17/2/503323 awarded to MN.

## References

1. Hu JX, Thomas CE, Brunak S. Network biology concepts in complex disease comorbidities. Nat Rev Genet. 2016;17(10):615–629.

2. Rajamanickam A, Munisankar S, Dolla C, Menon PA, Thiruvengadam K, Nutman TB, et al. Helminth infection modulates systemic pro-inflammatory cytokines and chemokines implicated in type 2 diabetes mellitus pathogenesis. PLoS Negl Trop Dis. 2020;14(3):e0008101.

3. Jaccard J, Turrisi R. Interaction effects in multiple regression. 72. Sage; 2003.

4. Jaccard J, Wan CK, Turrisi R. The detection and interpretation of interaction effects between continuous variables in multiple regression. Multivariate Behav Res. 1990;25(4):467–478.

5. Smith EN, Kruglyak L. Gene-environment interaction in yeast gene expression. PLoS Biol. 2008;6(4):e83.

6. Rajamanickam A, Munisankar S, Bhootra Y, Dolla C, Thiruvengadam K, Nutman TB, et al. Metabolic consequences of concomitant strongyloides stercoralis infection in patients with type 2 diabetes Mellitus. Clin Infect Dis. 2019;69(4):697–704.

7. Jourdan PM, Lamberton PHL, Fenwick A, Addiss DG. Soil-transmitted helminth infections. Lancet. 2018;391(10117):252–265.

8. Yazdanbakhsh M, Kremsner PG, van Ree R. Allergy, parasites, and the hygiene hypothesis. Science. 2002;296(5567):490–494.

9. DeFronzo RA, Ferrannini E, Groop L, Henry RR, Herman WH, Holst JJ, et al. Type 2 diabetes mellitus. Nat Rev Dis Primers. 2015;1:15019.

10. Guest, B C, Park MJ, Johnson DR, Freund GG. The implication of proinflammatory cytokines in type 2 diabetes. Frontiers in Bioscience-Landmark. 2008;13(13):5187–5194.

11. Keogh RA, Doran KS. Group B Streptococcus and diabetes: Finding the sweet spot. PLoS Pathog. 2023;19(2):e1011133.

12. Yazdanbakhsh M, van den Biggelaar A, Maizels RM. Th2 responses without atopy: immunoregulation in chronic helminth infections and reduced allergic disease. Trends Immunol. 2001;22(7):372–377.

13. Nutman TB. Looking beyond the induction of Th2 responses to explain immunomodulation by helminths. Parasite Immunol. 2015;37(6):304–313.

14. Mabbott NA. The influence of parasite infections on host immunity to co-infection with other pathogens. Front Immunol. 2018;9:2579.

15. Berbudi A, Ajendra J, Wardani AP, Hoerauf A, bner MP. Parasitic helminths and their beneficial impact on type 1 and type 2 diabetes. Diabetes Metab Res Rev. 2016;32(3):238–250.

16. Camaya I, O’Brien B, Donnelly S. How do parasitic worms prevent diabetes? An exploration of their influence on macrophage and *β*-cell crosstalk. Frontiers in Endocrinology. 2023;14:1205219.

17. Greaves D, Coggle S, Pollard C, Aliyu SH, Moore EM. Strongyloides stercoralis infection. BMJ. 2013;347:f4610.

18. Ryan SM, Eichenberger RM, Ruscher R, Giacomin PR, Loukas A. Harnessing helminth-driven immunoregulation in the search for novel therapeutic modalities. PLoS Pathog. 2020;16(5):e1008508.

19. Chen B, Khodadoust MS, Liu CL, Newman AM, Alizadeh AA. Profiling tumor infiltrating immune cells with CIBERSORT. Methods Mol Biol. 2018;1711:243–259.

20. Turner JD, Pionnier N, Furlong-Silva J, Sjoberg H, Cross S, Halliday A, et al. Interleukin-4 activated macrophages mediate immunity to filarial helminth infection by sustaining CCR3-dependent eosinophilia. PLoS Pathog. 2018;14(3):e1006949.

21. Girgis NM, Gundra UM, Ward LN, Cabrera M, Frevert U, Loke P. Ly6C(high) monocytes become alternatively activated macrophages in schistosome granulomas with help from CD4+ cells. PLoS Pathog. 2014;10(6):e1004080.

22. Sun L, Ye RD. Role of G protein-coupled receptors in inflammation. Acta Pharmacol Sin. 2012;33(3):342–350.

23. Kern PA, Ranganathan S, Li C, Wood L, Ranganathan G. Adipose tissue tumor necrosis factor and interleukin-6 expression in human obesity and insulin resistance. Am J Physiol Endocrinol Metab. 2001;280(5):E745–751.

24. Babu S, Kumaraswami V, Nutman TB. Alternatively activated and immunoregulatory monocytes in human filarial infections. The Journal of Infectious Diseases. 2009;199(12):1827–1837.

25. Rajamanickam A, Munisankar S, Thiruvengadam K, Menon PA, Dolla C, Nutman TB, et al. Impact of helminth infection on metabolic and immune homeostasis in non-diabetic obesity. Front Immunol. 2020;11:2195.

26. Rennie C, Fernandez R, Donnelly S, McGrath KC. The impact of helminth infection on the incidence of metabolic syndrome: A systematic review and meta-analysis. Front Endocrinol (Lausanne). 2021;12:728396.

27. Ayelign B, Negash M, Andualem H, Wondemagegn T, Kassa E, Shibabaw T, et al. Association of IL-10 (- 1082 A/G) and IL-6 (- 174 G/C) gene polymorphism with type 2 diabetes mellitus in Ethiopia population. BMC Endocr Disord. 2021;21(1):70.

28. Tuttle HA, Davis-Gorman G, Goldman S, Copeland JG, McDonagh PF. Proinflammatory cytokines are increased in type 2 diabetic women with cardiovascular disease. J Diabetes Complications. 2004;18(6):343–351.

29. Klotz C, Ziegler T, Figueiredo AS, Rausch S, Hepworth MR, Obsivac N, et al. A helminth immunomodulator exploits host signaling events to regulate cytokine production in macrophages. PLoS Pathog. 2011;7(1):e1001248.

30. Aira N, Andersson AM, Singh SK, McKay DM, Blomgran R. Species dependent impact of helminth-derived antigens on human macrophages infected with Mycobacterium tuberculosis: Direct effect on the innate anti-mycobacterial response. PLoS Negl Trop Dis. 2017;11(2):e0005390.

31. Huang H, Dong X, Kang MX, Xu B, Chen Y, Zhang B, et al. Novel blood biomarkers of pancreatic cancer–associated diabetes mellitus identified by peripheral blood–based gene expression profiles. Official journal of the American College of Gastroenterology— ACG. 2010;105(7):1661–1669.

32. Prada-Medina CA, Fukutani KF, Pavan Kumar N, Gil-Santana L, Babu S, Lichtenstein F, et al. Systems immunology of diabetes-tuberculosis comorbidity reveals signatures of disease complications. Scientific reports. 2017;7(1):1999.

33. Schmid DW, Fackelmann G, Wasimuddin, Rakotondranary J, Ratovonamana YR, Montero BK, et al. A framework for testing the impact of co-infections on host gut microbiomes. Animal microbiome. 2022;4(1):48.

34. Seber GA, Lee AJ. Linear regression analysis. vol. 330. John Wiley & Sons; 2003.

35. Wang X, Elston RC, Zhu X. The meaning of interaction. Hum Hered. 2010;70(4):269–277.

36. Hamilton NE, Ferry M. ggtern: Ternary diagrams using ggplot2. Journal of Statistical Software. 2018;87:1–17.

37. Jiang P, Zhang Y, Ru B, Yang Y, Vu T, Paul R, et al. Systematic investigation of cytokine signaling activity at the tissue and single-cell levels. Nat Methods. 2021;18(10):1181–1191.

38. Liao Y, Wang J, Jaehnig EJ, Shi Z, Zhang B. WebGestalt 2019: gene set analysis toolkit with revamped UIs and APIs. Nucleic Acids Res. 2019;47(W1):W199–W205.

39. Scutari M. Learning Bayesian networks with the bnlearn R package. arXiv preprint arXiv:09083817. 2009;.

40. Yuen KK. The two-sample trimmed t for unequal population variances. Biometrika. 1974;61(1):165–170.

