## Supplemental Information for "Delineating markers of disease-disease interaction: a systematic methodology and its application to multiple diabetes-helminth cohorts"

### 1 Supplementary Table

| Samples | Covariates | | | | | |
|---|---|---|---|---|---|---|
| | Age | ALT | AST | BMI | Creatinine | Sex (number of females/ number of males) |
| DM- Control | 39.02±11.85 | 20.35±13.98 | 21.85±7.61 | 21.95±1.54 | 0.79±0.19 | 25/35 |
| DM+ Control | 43.91±11.01 | 20.41±14.21 | 22.45±7.71 | 23.87±2.76 | 0.79±0.19 | 25/33 |
| DM- Pre-Treatment | 43.23±12 | 20.35±13.98 | 22.33±7.61 | 21.92±1.7 | 0.79±0.19 | 27/33 |
| DM+ Pre-Treatment | 45.33±11.34 | 25.95±16.96 | 24.35±13.37 | 20.51±2.95 | 0.80±0.22 | 30/30 |
| DM- Post-Treatment | 35.25±9.76 | 19.59±10.55 | 25.41±14.4 | 26.9±5.67 | 0.74±0.15 | 19/25 |
| DM+ Post-Treatment | 45.33±11.34 | 28.63±24.8 | 26.63±15.44 | 23.22±2.49 | 0.69±0.14 | 30/30 |

Table S1: **Distribution of covariates in the six cohorts.** This data file (.xlsx) lists the mean±sd (standard deviation) of continuous covariates like Age, AST, ALT, BMI, and Creatinine, and the number of occurrences of each value that a discrete covariate like Sex takes, in the six cohorts analyzed in this study.

### 2 Supplementary Figures

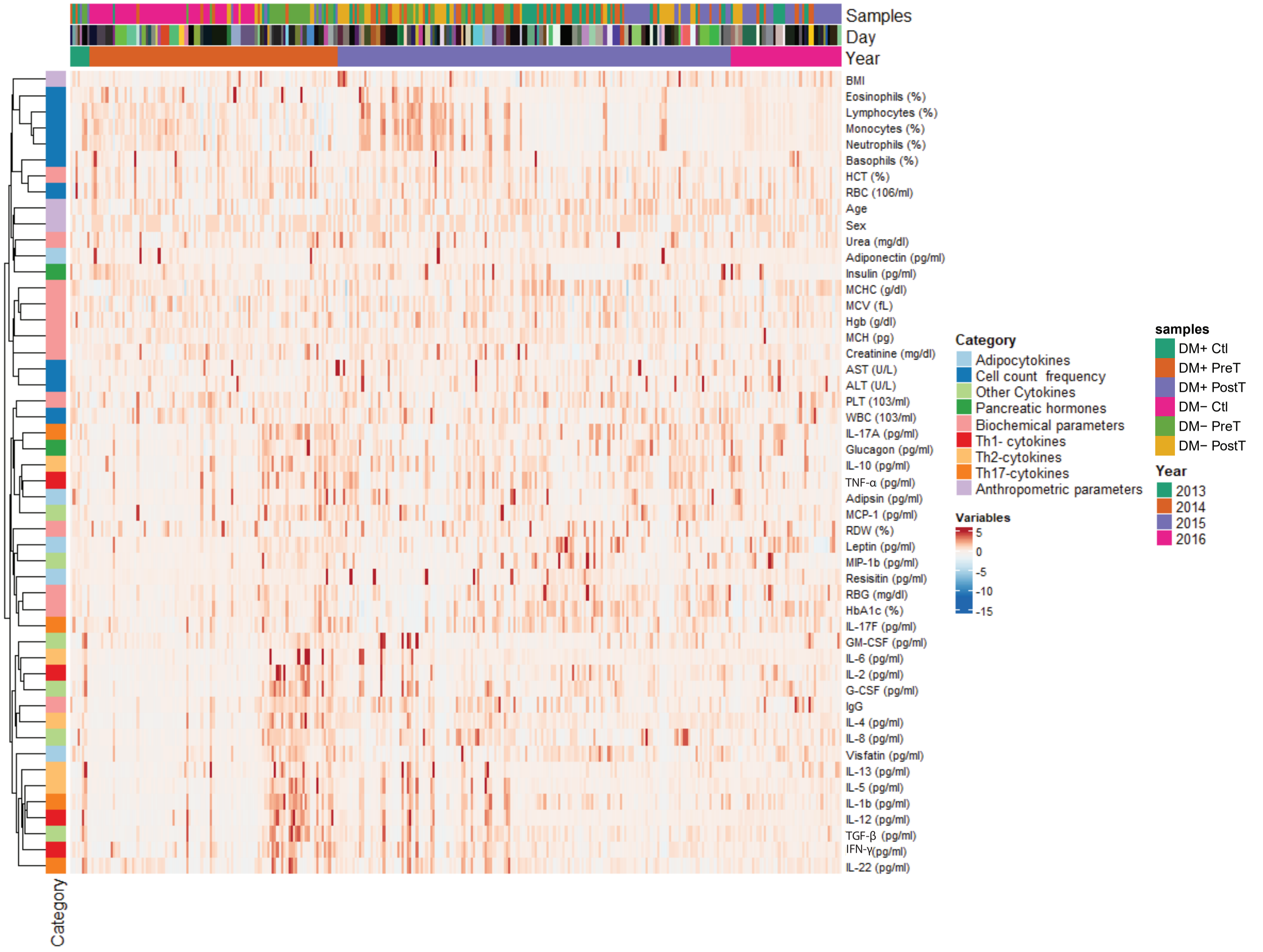

**Fig. S1. Overview of adjusted data:**. Heatmap of covariate-adjusted dataset which was obtained by concatenating data from six cohorts related to diabetes/helminth and adjusting the measured variables for these covariates: Age, Sex, Year, BMI, Creatinine, AST, and ALT (for reference, these covariates themselves are displayed as is, without adjustment, in this heatmap). Here “Day” refers to the date on which the samples were collected. In total, there are 171 unique dates at which the sample collection occurred from the year 2013 to 2016.

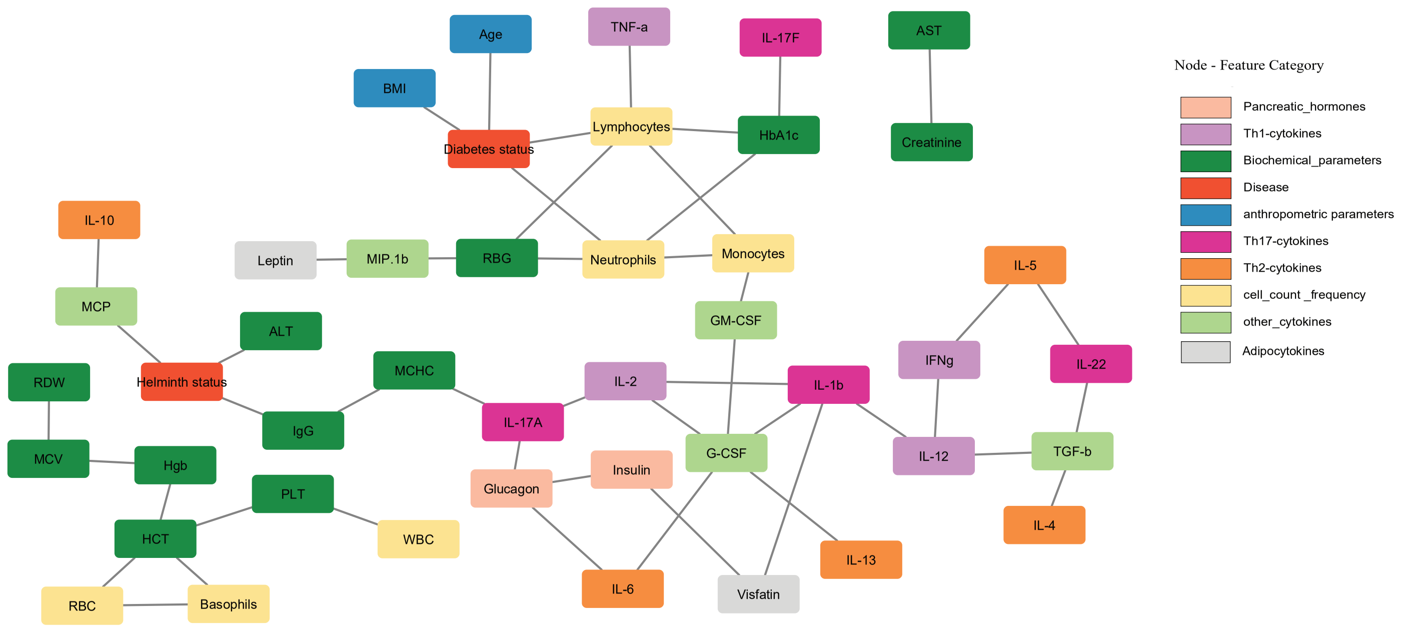

**Fig. S2. Before-treatment data:** Structure learning of the Bayesian network was employed to reconstruct a graph among the markers measured in the Control and Pre-treatment samples. Two disease nodes Diabetes status and Helminth status were introduced into the network to see how they affect the overall network modules. The variables measured have been color-coded based on the category they belong to (see S1 Table). The disease status nodes were encoded using the IgG level for helminth and HbA1c/RBG levels for diabetes (see main text, Results section).

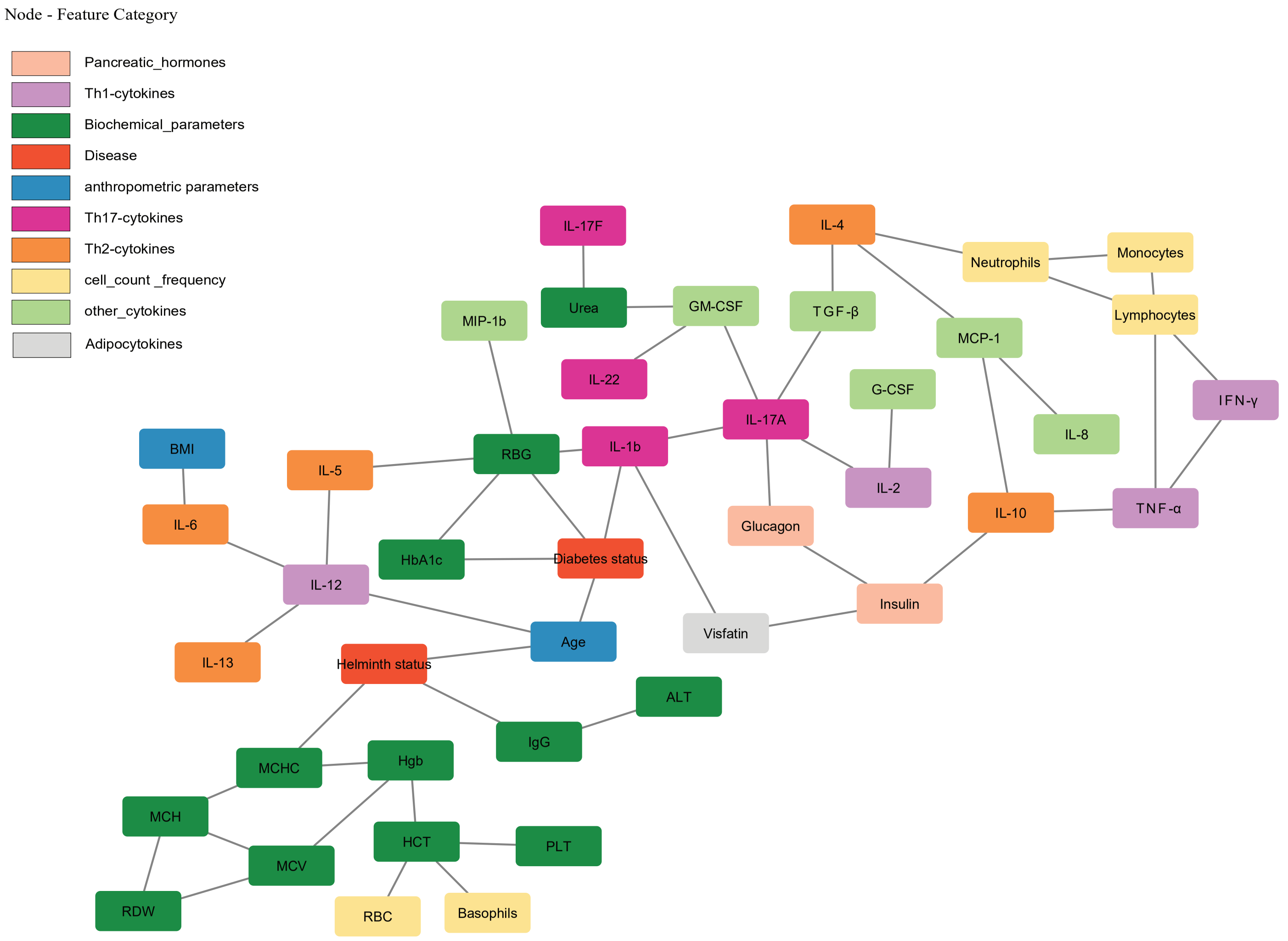

**Fig. S3. After-treatment data:** Structure learning of the Bayesian network was employed to reconstruct a graph among the markers measured in the Control and Post-treatment samples. Two disease nodes Diabetes status and Helminth status were introduced into the network to see how they affect the overall network modules. The variables measured have been color-coded based on the category they belong to (see S1 Table). The disease status nodes were encoded using the IgG level for helminth and HbA1c/RBG levels for diabetes (see main text, Results section).

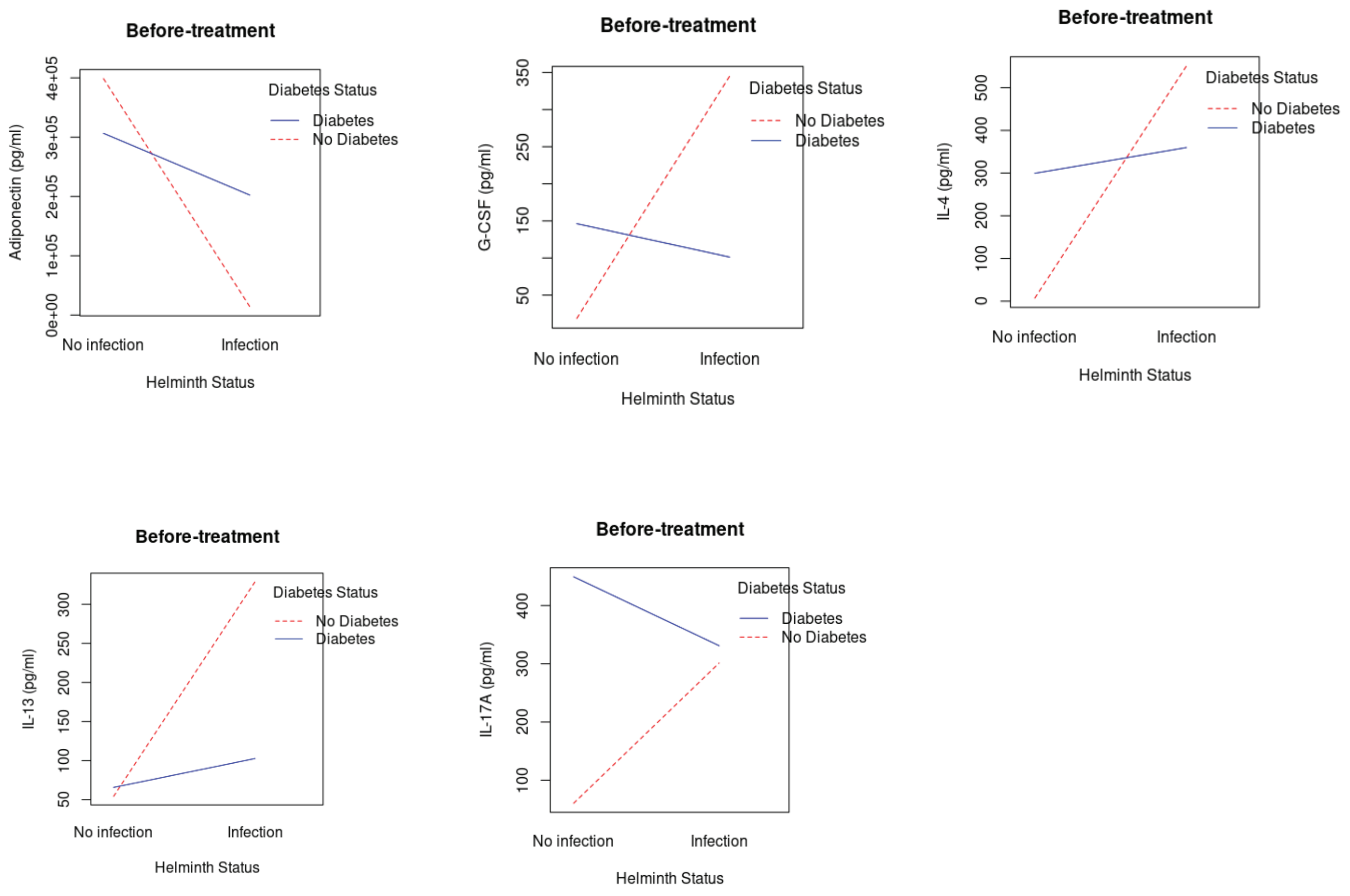

**Fig. S4. Interaction plots (before-treatment):** Interaction plots for the variables like Adiponectin, IL-13, IL-4, G-CSF, and IFN-$\gamma$ that are mentioned in Fig 3A for the before-treatment samples are provided here. Here, the x-axis represents the status of helminth infection and the lines (red-dotted and blue-straight) represent the status of diabetes.

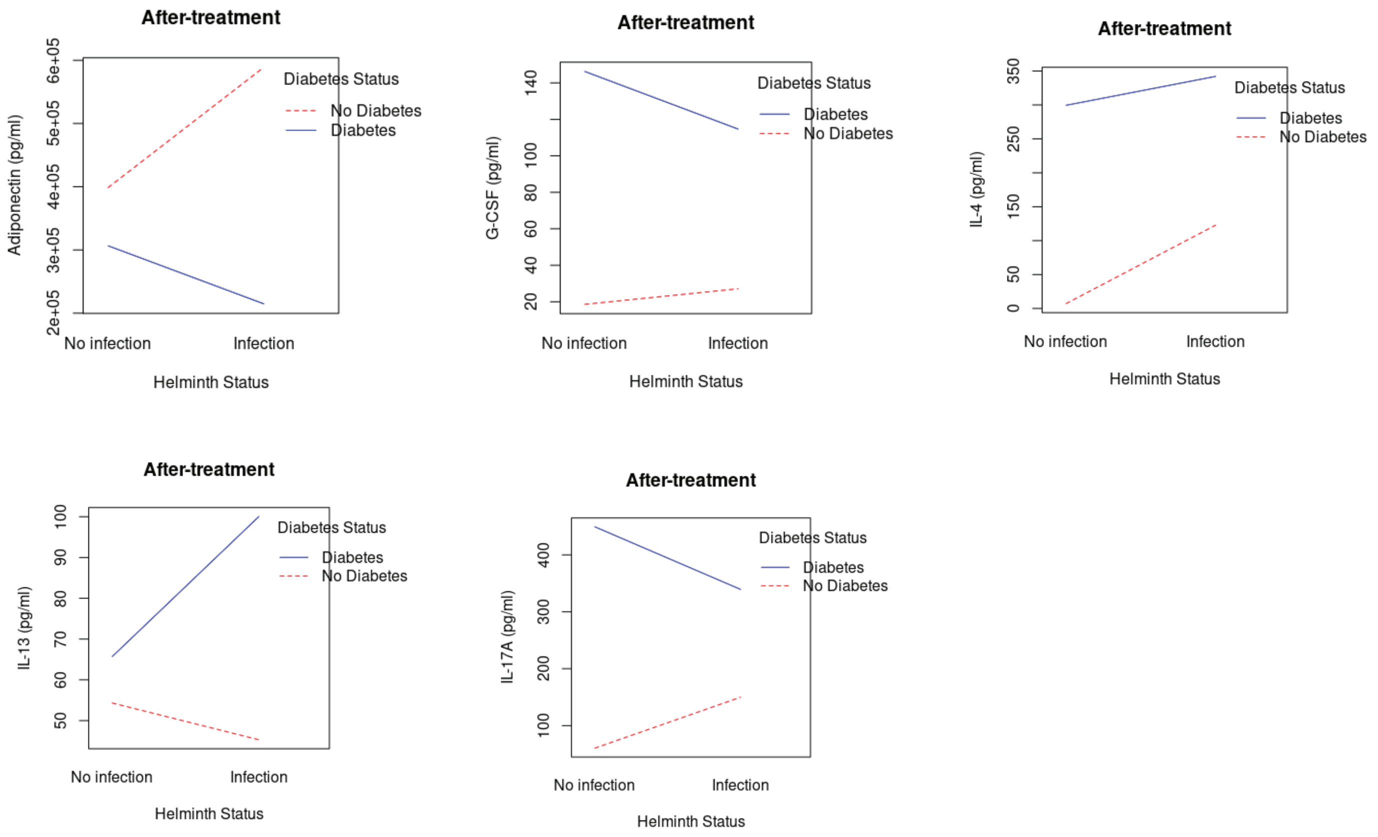

**Fig. S5. Interaction plots (after-treatment):** Interaction plots for the variables like Adiponectin, IL-13, IL-4, G-CSF, and IFN-gamma that are mentioned in Fig 3B for the after-treatment samples are provided here. Here, the x-axis represents the status of helminth infection and the lines (red-dotted and blue-straight) represent the status of diabetes.

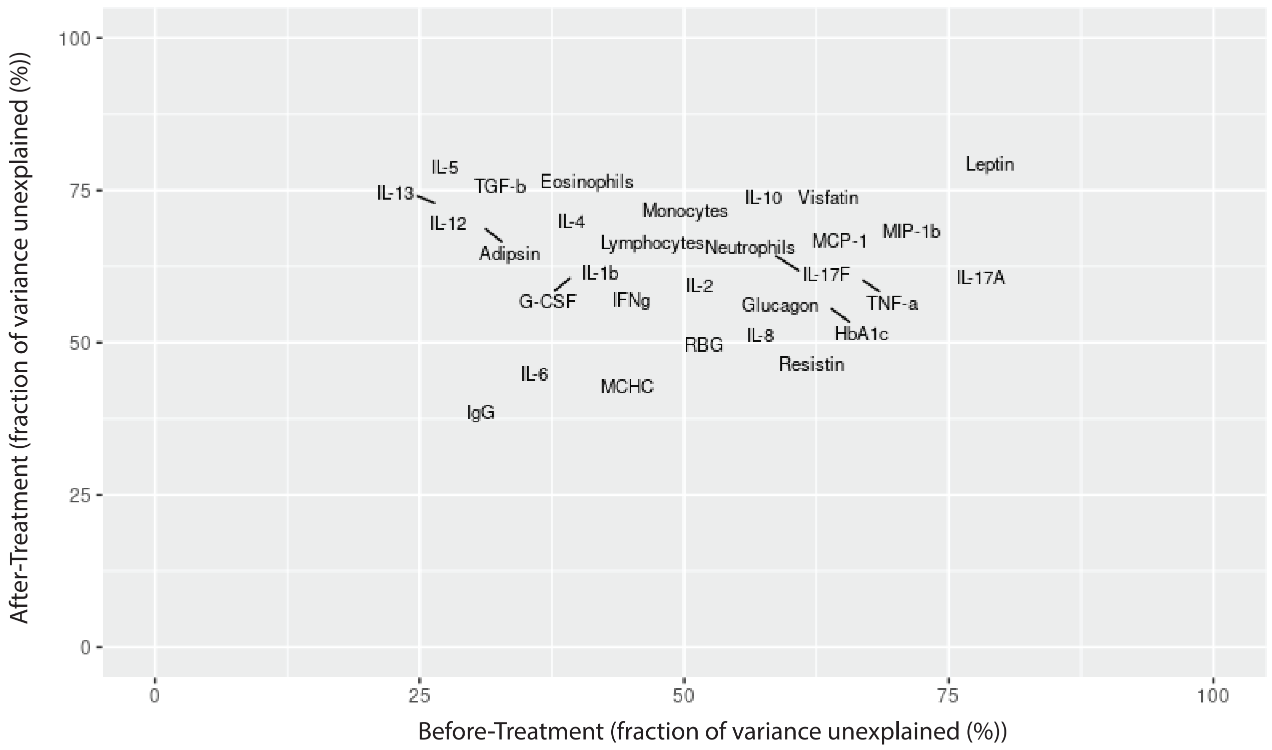

**Fig. S6. Unexplained variance plot:** Before vs. after anthelmintic treatment plot of the percentage of marker variance that is unexplained for the main effect markers (markers that had a significant main effect component in before- or after-treatment data; see Methods). The values used for this plot are given in S2 and S3 Tables.

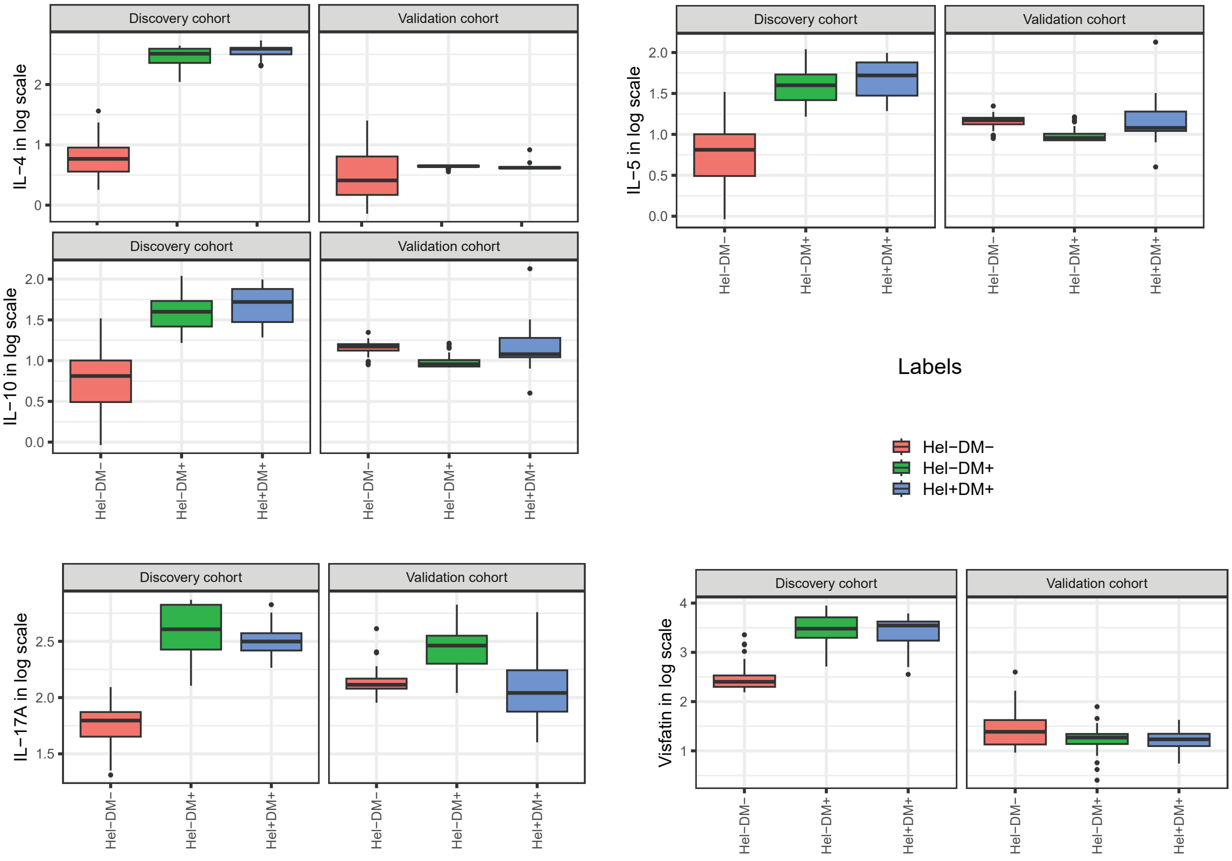

**Fig. S7. Additional replication results:** Expression levels of certain Th2 cytokines (IL-4, IL-5, and IL-10), Th17 cytokine IL-17A, and Visfatin are shown after (base-10) log transformation in both discovery and validation cohorts.

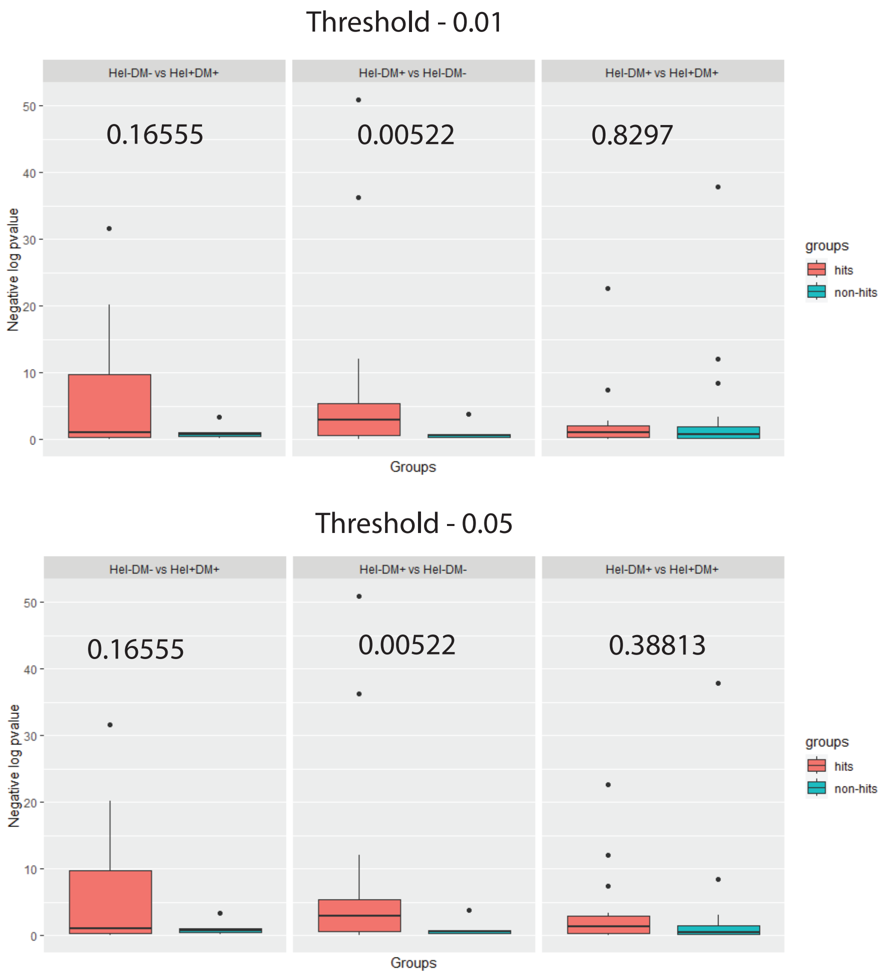

**Fig. S8. DE-based replication testing:** DE signals ($-log_{10}$(DE p-value)) in the validation dataset of the DE hits vs. non-hits that were called in the discovery dataset (at an adjusted p-value threshold of 0.01 and 0.05). Whether DE signals of hits are better than that of non-hits was tested using Yuen's robust t-test and its p-value is shown for each comparison.

### 3 Supplementary Files

Supplementary data files listed below are available at this link: https://github.com/BIRDSgroup/DDI/tree/main/Application%20on%20helminth-diabetes%20data/Supplementary%20File%20and%20Data.

Suppl File D1: **Measured variables and their category.** 50 variables are measured from various cohorts, and they are categorized based on their origin (cells that produce them), biochemical parameters (hormones, liver and excretory measurements, RBC parameters, antibody produced), and anthropometric parameters.

Suppl File D2: **Main and Interaction effects - Before treatment.** This table contains the following information from fitting linear regression models to the before-treatment data: the considered variable, main and interaction effect p-values (before and after multiple testing adjustment), coefficients from the linear model in Eq 2 (Intercept term, Helminth, and Diabetes; refer to manuscript), the relative proportion of variance explained by helminth, diabetes and interaction term, fraction of variance unexplained (by these three terms), and the coefficients from the linear model in Eq 3 (Intercept term, Helminth, Diabetes and Helminth:Diabetes interaction term; refer to manuscript).

Suppl File D3: **Main and Interaction effects - After treatment.** This table consists of the same information as S2 Data, but with the linear regression models fit to the after-treatment data.

Suppl File D4: **Replication analysis - hits and non-hits.** This table lists the hits and non-hits called in the discovery cohort at various FDR cut-offs (0.0001, 0.01 and 0.05).

Suppl File D5: **Top 100 entries of TGs.** This data file (.xlsx) consists of the top 100 entries (search results) when Cytosig is queried/searched for the target genes of certain DDI markers of interest.

Suppl File D6: **Enrichment p-value of Reactome pathways for cytokine TGs.** This data file (.csv) lists the p-values corresponding to enrichment of the Reactome pathways for the top TGs (target genes) of certain DDI cytokines.
